# Small molecule structure elucidation from NMR spectra with generative and reasoning models

**DOI:** 10.64898/2026.10.29.762711

**Authors:** Ziyu Xiong, Vanessa Y. Ying, Jonathan Z. Huang, Zane Koch, Joseph D. Clark, Robert C. Heeter, Joon Soo An, Karit Matanachai, Samuel G. Rodriques, Andrew D. White, Mohammad R. Seyedsayamdost, Ellen D. Zhong

**Author notes:** Corresponding author(s): Ellen D. Zhong and Mohammad R. Seyedsayamdost.

## Abstract

Elucidating the structures of unknown small molecules remains a major challenge in chemistry, pharmacology, and drug discovery. While nuclear magnetic resonance (NMR) spectroscopy is the gold standard for structure elucidation, interpreting spectra remains labor-intensive and requires expert knowledge to navigate the vast space of possible structures. Here, we present an AI system for rapid, automated *de novo* structure elucidation that combines a molecular generative model with a scientific reasoning agent. Its generative component, ChefNMR2, is an atomic flow-matching model that predicts small-molecule structures from molecular formulae and one-dimensional ^1^H and ^13^C NMR spectra. ChefNMR2 - Agent augments our generative model with large language models (LLMs) to reason over and refine candidate structures. Across diverse benchmarks, ChefNMR2 substantially outperforms existing tools, offering calibrated confidence estimates and two to three orders of magnitude faster inference than frontier LLMs. We use ChefNMR2 - Agent to elucidate the structures of four novel natural products, matching human-chemist accuracy in a fraction of the time. Our work demonstrates how specialized generative models and general-purpose reasoning agents can unite to solve challenging scientific inverse problems and accelerate NMR structure elucidation, thus opening new possibilities for AI-driven small-molecule discovery across diverse disciplines.

## 1. Introduction

Naturally occurring small molecules have revolutionized science and medicine and provided some of our most celebrated cures. They account for more than half of all FDA-approved drugs and remain a steady source for therapeutic agents ^1^.However, a major bottleneck in natural product and small-molecule discovery is the structure-elucidation paradox ^2,3^. While bioactive hits can be easily identified, determining structures *de novo* is a slow and painstaking endeavor ^2^. Nuclear magnetic resonance (NMR) spectroscopy is the principal analytical technique for this process ^4,5^ (Fig. 1a). But inferring structures from NMR data remains a largely manual exercise that requires substantial time, expert interpretation, and long spectral acquisition of two-dimensional (2D) NMR data that provide information on nuclear connectivity ^5–9^. Automating structure elucidation from simple one-dimensional (1D) NMR spectra alone remains an open research problem, with the potential to substantially expedite small-molecule characterization and discovery.

**Fig. 1.**
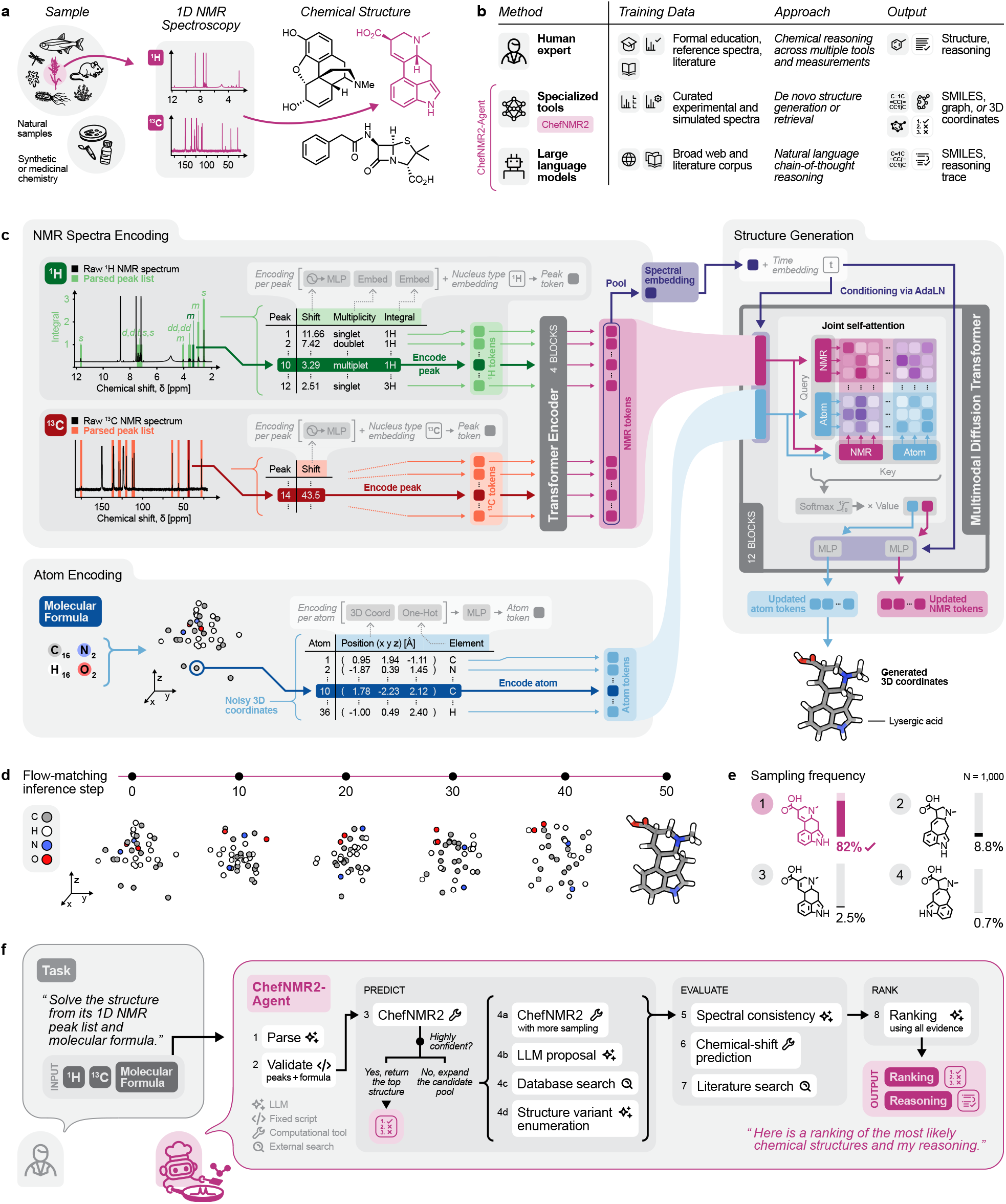
Automated NMR structure elucidation with ChefNMR2 and ChefNMR2 - Agent. **a**, The task of molecular structure elucidation from one-dimensional nuclear magnetic resonance (NMR) spectra. **b**, Overview of approaches to structure elucidation, including human experts, specialized tools and large language models (LLMs). SMILES, simplified molecular-input line-entry system. **c**, ChefNMR2 architecture, comprising a peak encoder and a multimodal diffusion transformer with joint attention between peak and atom tokens. AdaLN, adaptive layer normalization. MLP, multilayer perceptron. *t*, flow-matching time. **d**, An example flow-matching trajectory from Gaussian noise to atomic coordinates of lysergic acid. Colors denote elements. **e**, Candidate structures ranked by sampling frequency, with the correct structure ranked first (pink). *N*, number of samples drawn. **f**, Workflow of ChefNMR2 - Agent, integrating ChefNMR2 with chemical reasoning and computational tools for candidate prediction, evaluation and ranking.

Computational methods have long sought to automate structure elucidation ^7^. Classical computer-assisted structure elucidation (CASE) programs derive structures consistent with NMR-based constraints but rely primarily on 2D NMR correlations ^8^. More recently, deep learning approaches have been developed to solve *de novo* structures ^6,10–18^, or retrieve candidates from a predefined library ^15,19,20^. However, current methods perform poorly on experimental spectra, thus limiting their practical use ^6,19,21,22^. Nascent advances in large language models (LLMs) offer another paradigm for computational structure elucidation (Fig. 1b). Beyond their broad chemical knowledge, LLMs demonstrate emerging capabilities in chemical reasoning, prediction of reaction outcomes, and planning synthetic schemes ^23^. But although general-purpose LLMs can reason over molecular formulae and spectral data to propose candidate structures ^24,25^, their accuracy and reliability remain poor, particularly for complex molecules and natural products ^26,27^. Beyond standalone reasoning, LLMs can also function as scientific agents, orchestrating external computational tools to solve complex scientific problems ^28–30^. Nonetheless, the effectiveness of integrating LLM agents with specialized models for structure elucidation remains underexplored ^27,31,32^.

Here we introduce ChefNMR2 and ChefNMR2 - Agent to combine molecular generation with chemical reasoning agents for automated structure elucidation from 1D NMR spectra (Fig. 1c– f). ChefNMR2 is an atomic flow-matching model trained on 2.52 million small molecules with experimental and simulated spectra to predict structures from molecular formulae and ^1^H and ^13^C NMR peak lists. We benchmark ChefNMR2 against both extant computational methods and frontier LLMs and find that it substantially outperforms all existing methods, with structure elucidation rates 2–3 orders of magnitude faster than frontier LLMs. Moreover, ChefNMR2 provides calibrated confidence estimates that we use to guide subsequent analysis by an LLM-based scientific agent. In our integrated system, ChefNMR2 - Agent evaluates and refines ChefNMR2 predictions through complementary chemical tools and web search, guided by the chemical reasoning capabilities of general-purpose LLMs. We use this system to solve the structures of four novel natural products, including those that elude both the generative model and standalone LLM reasoning, achieving human-level accuracy in a fraction of the time and without 2D NMR spectra. These results establish the combination of specialized generative models with scientific agents to solve challenging problems at the chemistry-life science interface. Both ChefNMR2 and ChefNMR2 - Agent are openly available.

## 2. Results

### 2.1. Overview of ChefNMR2 and ChefNMR2 - Agent

ChefNMR2 is a small-molecule generative model trained to capture the complexity and diversity of chemical space for *de novo* structure elucidation. We frame structure elucidation as generating atomic coordinates conditioned on a molecular formula and one-dimensional ^1^H and ^13^C NMR peak lists. Inspired by multimodal diffusion transformers that jointly process image and text representations ^33^, we develop an architecture that jointly processes an atomic representation of the molecule and NMR spectral peaks. Within ChefNMR2, a molecule’s molecular formula defines the set of atoms, while an encoder processes input 1D NMR peak lists as spectral tokens. Joint attention between these token streams allows NMR spectral tokens to guide the evolving atomic coordinates, with a pooled spectral embedding providing additional conditioning through adaptive layer normalization (Methods 4.2.3, Fig. 1c).

We train the model using a flow-matching objective ^34^. Random rotation of molecular conformers during training encourages the model to learn rotational symmetry without enforcing equivariance in the architecture, similar to recent trends in the protein structure modeling literature ^35^. We greatly expand the training data relative to prior work ^18^ to 2.52 million molecules (Fig. 2a), combining experimental NMR data mined from the chemistry literature ^36^ with simulated spectra of synthetic compounds from US patents and natural products from COCONUT, NPAtlas and NP-MRD ^21,37–40^ (Methods 4.1).

**Fig. 2.**
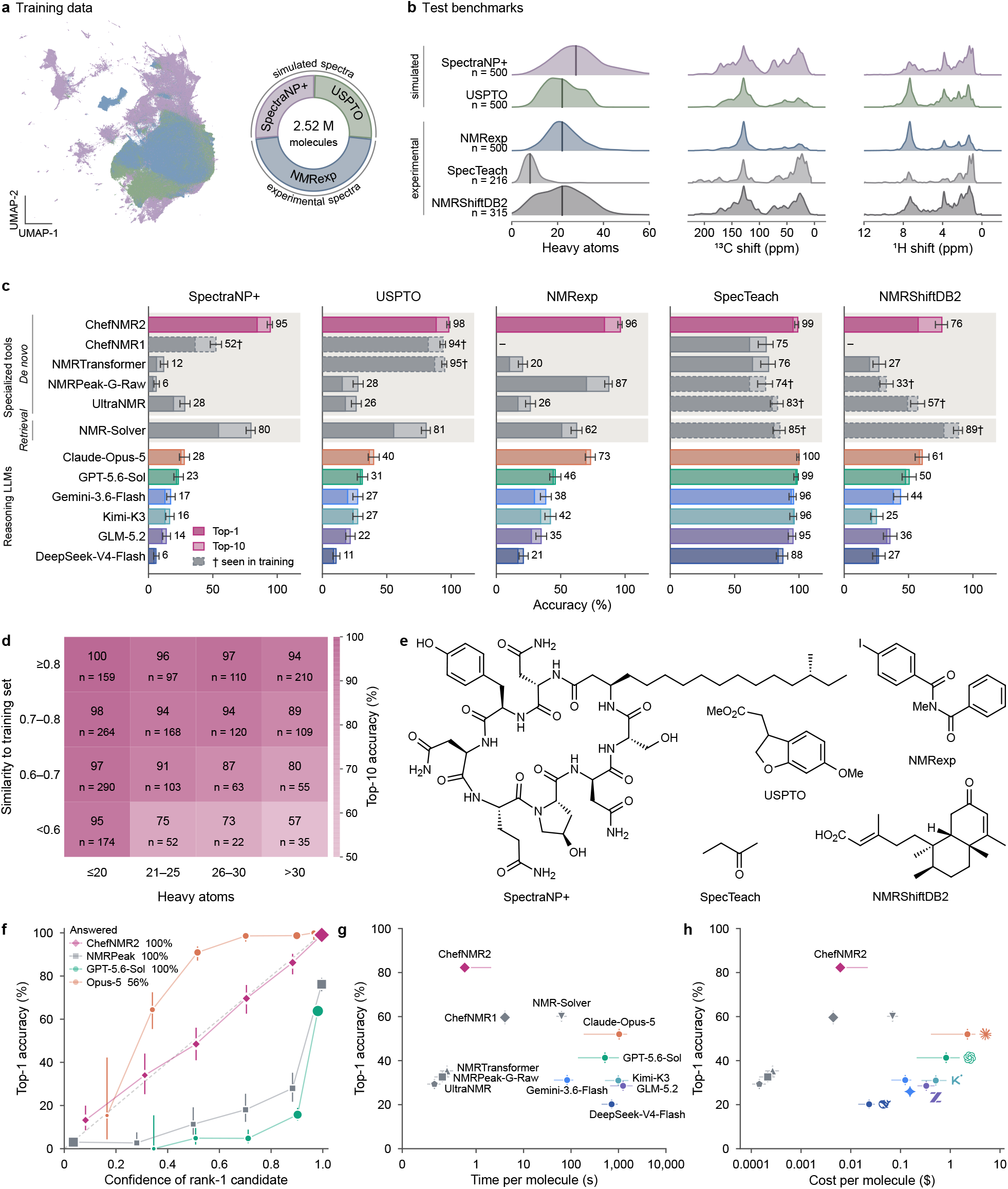
Accurate and calibrated structure elucidation by ChefNMR2. **a**, Left, uniform mani-fold approximation and projection (UMAP) of ChefNMR2’s spectral embeddings of the training molecules, colored by dataset. Right, share of molecules in each dataset. **b**, Distributions of heavy atoms per molecule (tick, median) and of ^13^C and ^1^H chemical shifts. *n*, number of molecules. **c**, Top-1 (dark) and top-10 (light) accuracy. Candidates are matched by planar structure. Numbers, top-10 accuracy. Set sizes as in **b**. Seen in training (†, dashed outline), at least 5% of the set is in that tool’s training data. –, not applicable. **d**, Top-10 accuracy of ChefNMR2 by number of heavy atoms and Tanimoto similarity to the nearest training molecule. **e**, Examples of correct top-1 predictions by ChefNMR2, one from each test set. **f**, Top-1 accuracy against the confidence of the rank-1 candidate for ChefNMR2 (sampling frequency), NMRPeak-G-Raw (probability) and two LLMs (stated confidence), over the molecules each system answers (percentages). Marker size increases with the number of molecules. Dashed line, confidence equal to accuracy. **g**,**h**, Top-1 accuracy against median runtime (**g**) and cost (**h**) per molecule. Horizontal bars, 25th to 75th percentiles. Error bars, 95% Wilson intervals (top-10 accuracy in **c**, top-1 elsewhere). In **d** and **f**–**h**, the test sets are pooled.

At inference time, the model transforms random noise into atomic coordinates, from which we assign bonds to obtain candidate structures (Fig. 1d). We estimate the uncertainty of the predictions by repeated sampling through the generative model, using the frequency of each distinct planar structure as an empirical confidence estimate (Fig. 1e).

While ChefNMR2 provides the core generative capability, recent advances in general-purpose LLMs and scientific agents offer complementary ways to interpret NMR spectra, drawing on chemical knowledge and reasoning much as a chemist would. We bring these capabilities together in ChefNMR2 - Agent, which provides a natural-language interface and extends molecular generation with chemical reasoning, computational tools, and literature search (Fig. 1f). The agent uses ChefNMR2’s confidence estimates to adapt the amount of sampling and analysis, returning confident predictions quickly and investigating uncertain cases further (Methods 4.6.1). For uncertain cases, the agent can expand the candidate pool through independent LLM proposals, compare predicted chemical shifts for candidate molecules with observed peaks using chemical-shift prediction tools such as CSP5^41^, and consult the literature to help distinguish competing structures. ChefNMR2 - Agent integrates this information to rank candidate structures and explain the chemical reasoning behind its ranking. The full workflow and associated tools are described in the Methods.

### 2.2. Accurate and calibrated small-molecule structure elucidation

A key component of ChefNMR2 is the substantially expanded scale and diversity of its training data. We assemble 2.52 million molecules by combining experimental NMR peak lists mined from the literature (NMRexp)^36^ with simulated spectra of compounds from US patents (USPTO)^21,37^ and natural product databases (SpectraNP+). We construct SpectraNP+ from COCONUT, NPAtlas, and NP-MRD ^38–40^, extending the natural product dataset introduced in earlier work ^18^. A UMAP visualization of ChefNMR2’s learned spectral embeddings reveals overlapping and distinct regions occupied by the three training datasets (Fig. 2a). Analysis of their structural composition shows that SpectraNP+ is enriched in fused, bridged, spiro, aliphatic, and macrocyclic ring systems, whereas nitrogen-containing compounds and aromatic rings are more prevalent in USPTO and NMRexp (Fig. S1a,b).

We evaluate ChefNMR2 on a total of 2,031 molecules across five test sets (Fig. 2b). Three are held out from NMRexp, USPTO, and SpectraNP+, and we curate two additional test sets from public experimental NMR collections. NMRShiftDB2 provides assigned experimental peak lists ^42,43^ (Methods 4.3.1), while SpecTeach contains spectra for simple compounds collected for undergraduate teaching ^44^, from which we extract peak lists by automated processing (Methods 4.3.1).

We compare ChefNMR2 with five specialized structure elucidation tools and six general-purpose reasoning LLMs (Fig. 2c). The specialized tools include our earlier atomic generative model Chef NMR1^18^, three chemical language models that generate SMILES strings ^11,13,15^, and the retrieval method NMR-Solver ^19^. The LLMs are given the molecular formula and NMR peak lists and rely on their own chemical knowledge and reasoning, without external tools or web access. Across the five test sets, ChefNMR2 ranks the correct planar structure first for 82.3% of molecules, compared with 35.4% for the best specialized *de novo* model, NMRTransformer, 60.1% for the retrieval method NMR-Solver, and 52.0% for the best LLM, Claude-Opus-5. As some specialized baselines were trained on molecules in the test sets, we report the extent of this overlap and observe higher accuracy on seen than unseen molecules, consistent with inflated performance due to memorization (Fig. S2a).

We stratify ChefNMR2’s performance by size and similarity to the training data as a means to assess generalization (Fig. 2d). ChefNMR2 recovers the correct structure among its top ten candidates for 87.5% of molecules with more than 30 heavy atoms and 84.8% of molecules whose nearest training neighbor has a Tanimoto similarity below 0.6. Molecules that are both large and dissimilar remain more challenging, with top-10 accuracy of 57.1% across 35 compounds. Notably, compared with ChefNMR2, accuracy declines more sharply for the specialized baselines as similarity to their training data decreases, and for all specialized baselines and LLMs as molecular size increases (Fig. S2b,c). An example of correctly elucidated compounds by ChefNMR2 in each test set is shown in Fig. 2e.

An essential component of a structure prediction tool is a confidence metric with which it assigns structure. We compare ChefNMR2’s confidence estimates, measured by sampling frequency, with observed accuracy across the test sets (Fig. 2f). We find that the sampling frequency closely tracks accuracy, with an expected calibration error of 1.3 percentage points ^45^. By comparison, confidence estimates from the LLMs and three chemical language models deviate more substantially from their observed accuracy, with expected calibration errors of 12.6 to 49.4 percentage points (Fig. S2f). Thus, sampling frequencies provide an intrinsically calibrated confidence estimate, without the need for a separately trained confidence module.

ChefNMR2 uses prediction confidence to adapt its sampling budget after an initial 100 samples. Sampling stops when one structure accounts for at least 97% of the generated structures; otherwise, it expands to 1,000 or 3,000 samples depending on the confidence of the initial predictions (Methods 4.2.5). Across the five test sets, sampling terminates after 100 draws for 58.5% of molecules, and 99.2% of these predictions are correct. With this adaptive budget, ChefNMR2 achieves up to 2,170-fold lower median runtime and 360-fold lower median cost per molecule than the LLMs (Fig. 2g,h).

### 2.3. Elucidating complex natural products

Natural products provide the ultimate test for ChefNMR2’s ability to deduce complex molecular structures. We first evaluate nine well-known natural products spanning diverse structural classes not found within our experimental training data (Fig. 3a,b and Methods). ChefNMR2 is able to elucidate the correct planar structure for all nine at rank-1, including vancomycin aglycone, rapamycin and taxol, the largest and most complex examples in this set. In comparison, NMRTransformer ranks three correctly at the top, while NMRPeak-G-Raw identifies none.

**Fig. 3.**
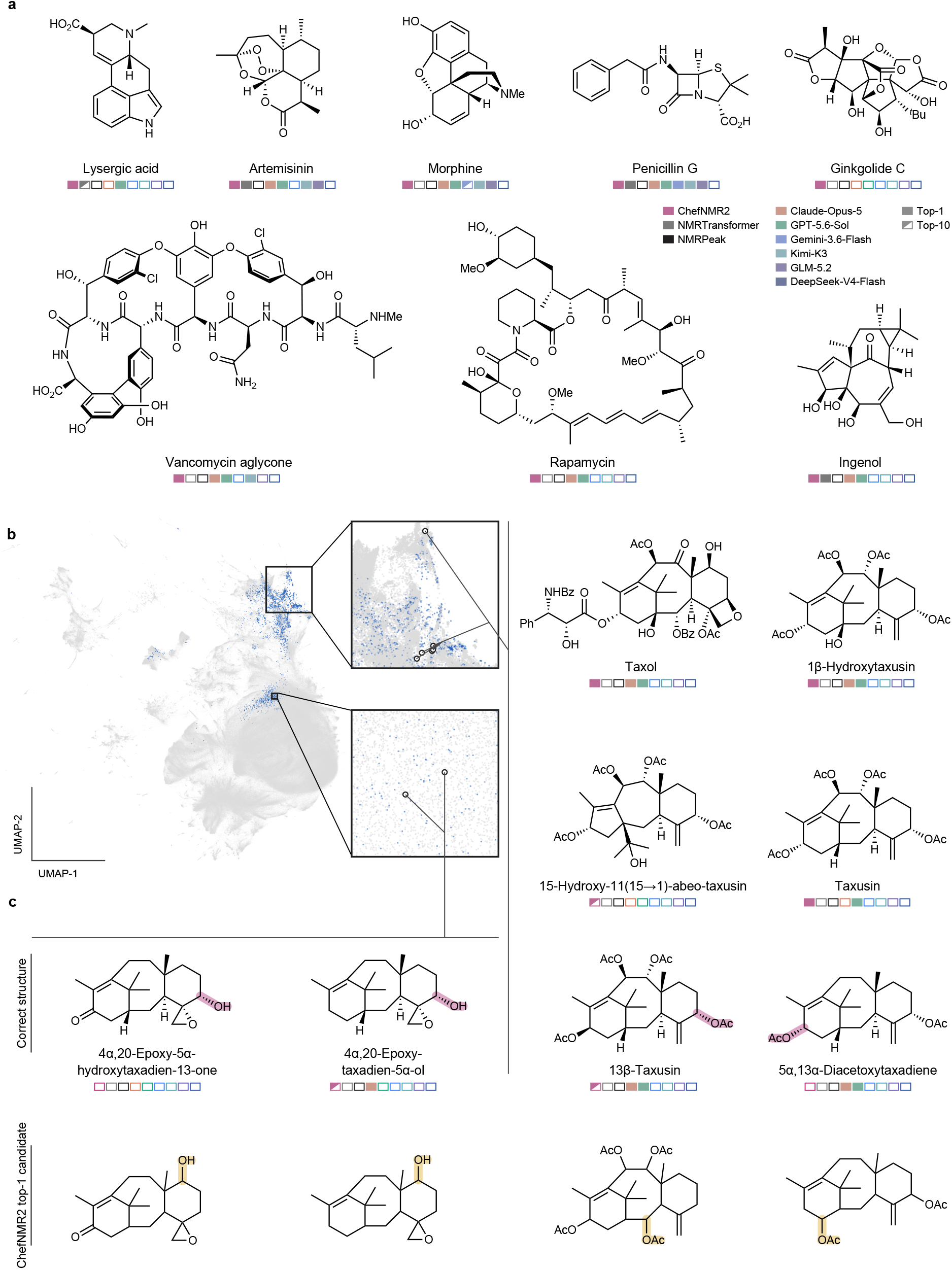
Identifying complex natural products and taxanes with ChefNMR2. **a**, Well-known natural products with diverse structures, none of which occurs in ChefNMR2’s experimental training data. Filled squares, correct structure ranked first. Half-filled squares, ranked second to tenth. Open squares, not in the top ten. Structures are matched by planar structure without consideration of stereochemistry. For ChefNMR2, a match requires at least three of five runs to rank the correct structure first (top-1) or within the top ten (top-10). All other systems are run once. NMR-Solver and UltraNMR are not shown because most compounds occur in their databases or training data; Chef NMR1 is omitted because it requires raw spectra. **b**, Left, uniform manifold approximation and projection (UMAP) of ChefNMR2’s spectral embeddings of the training molecules (gray). Blue, 1,000 training molecules nearest to each taxane in the embedding. Insets enlarge the regions that hold the five taxanes with acetoxy groups (top) and the two 4*α*,20-epoxides (bottom). **c**, Correct structures (top) and ChefNMR2’s top-1 candidates from one run (bottom) for four taxanes. Shading marks the groups that differ between each pair.

The frontier language models GPT-5.6-Sol and Claude-Opus-5 also perform well on these compounds, correctly identifying eight and seven, respectively. However, their responses suggest that the models have recognized these well-known molecules; each correct LLM response names the compound, and several begin by matching the molecular formula, e.g., to vancomycin aglycone or taxol (Table S1a,b). These observations make it difficult to distinguish structural inference from recall for well-known natural products.

We therefore examine seven closely related taxane diterpenoids from a study of taxol biosynthesis ^46^, testing whether the models can resolve structural differences within a shared molecular family (Fig. 3b,c). ChefNMR2 recovers the correct structure among its top ten candidates for five of the seven compounds. Even when its top-ranked prediction is incorrect, it can recover the ring system and hydroxy or acetoxy groups (Fig. 3c). NMRTransformer, NMRPeak-G-Raw, and four of the six LLMs identify none of the seven. GPT-5.6-Sol and Claude-Opus-5 each identify four, requiring 6–25 minutes per molecule. In contrast, Claude-Opus-5 identifies taxol in approximately one minute, further suggesting that familiarity contributes to its performance on the well-known compound (Table S1b).

The model’s learned spectral representations also reflect structural relationships among these molecules. In the UMAP projection, the five taxanes with acetoxy groups cluster near taxol, while the two 4*α*,20-epoxides occupy a separate region (Fig. 3b). More broadly, principal components of the learned spectral embeddings correlate with chemically interpretable properties, including molecular size and the fraction of sp^3^ carbons (Fig. S3d). These relationships suggest that ChefNMR2 learns a chemically meaningful latent space that could support downstream tasks beyond structure prediction, such as prediction of biochemical or pharmacological properties of test compounds.

### 2.4. Discovery of novel natural product structures with ChefNMR2 - Agent

As a final test, we apply ChefNMR2 and the LLM-based agent toward the discovery and structure elucidation of novel natural products. We selected *Nocardia transvalensis*, the causative agent of the infectious disease nocardiosis, and *Streptomyces* sp. HPH0547, a commensal microbiome-associated strain. Both microbes are rich in natural product biosynthetic pathways. Large-scale cultures allowed us to isolate two novel natural products from *N. transvalensis*. Their structures were elucidated by a human chemist using 1D/2D NMR spectra. Subsequently, the 1D spectra were supplied to ChefNMR2 - Agent.

For the first analog, ChefNMR2 predicts a largely correct structure but favors an isomer containing *α*-methoxyglycine and ranks the correct, serine-containing structure fourth or fifth (Fig. 4a). ChefNMR2 - Agent distinguishes these alternatives by examining the peaks associated with the bridging serine amino acid. Chemical-shift predictions conducted by the agent using CSP5^41^ place the ^13^C shift of serine’s CH_2_OH group close to the observed peak at 61.7 ppm; the agent assigns the ^1^H peak at 3.70 ppm to the same group. The competing isomer leaves these signals unexplained. Literature searches ^47–49^ further identify serine in a related siderophore, gobichelin B ^50^. Together, these observations lead the agent to rank the correct structure first in three of five runs. Without ChefNMR2, the agent usually fails to identify it. Moreover, all specialized baselines and LLMs tested fail to predict the correct structure. Because ChefNMR2 - Agent recovers this structure from 1D spectra alone, we name it nocobactin AI-1. Interestingly, it is a member of the nocobactins, which have been suggested as virulence factors of pathogenic *Nocardia* species ^51^.

**Fig. 4.**
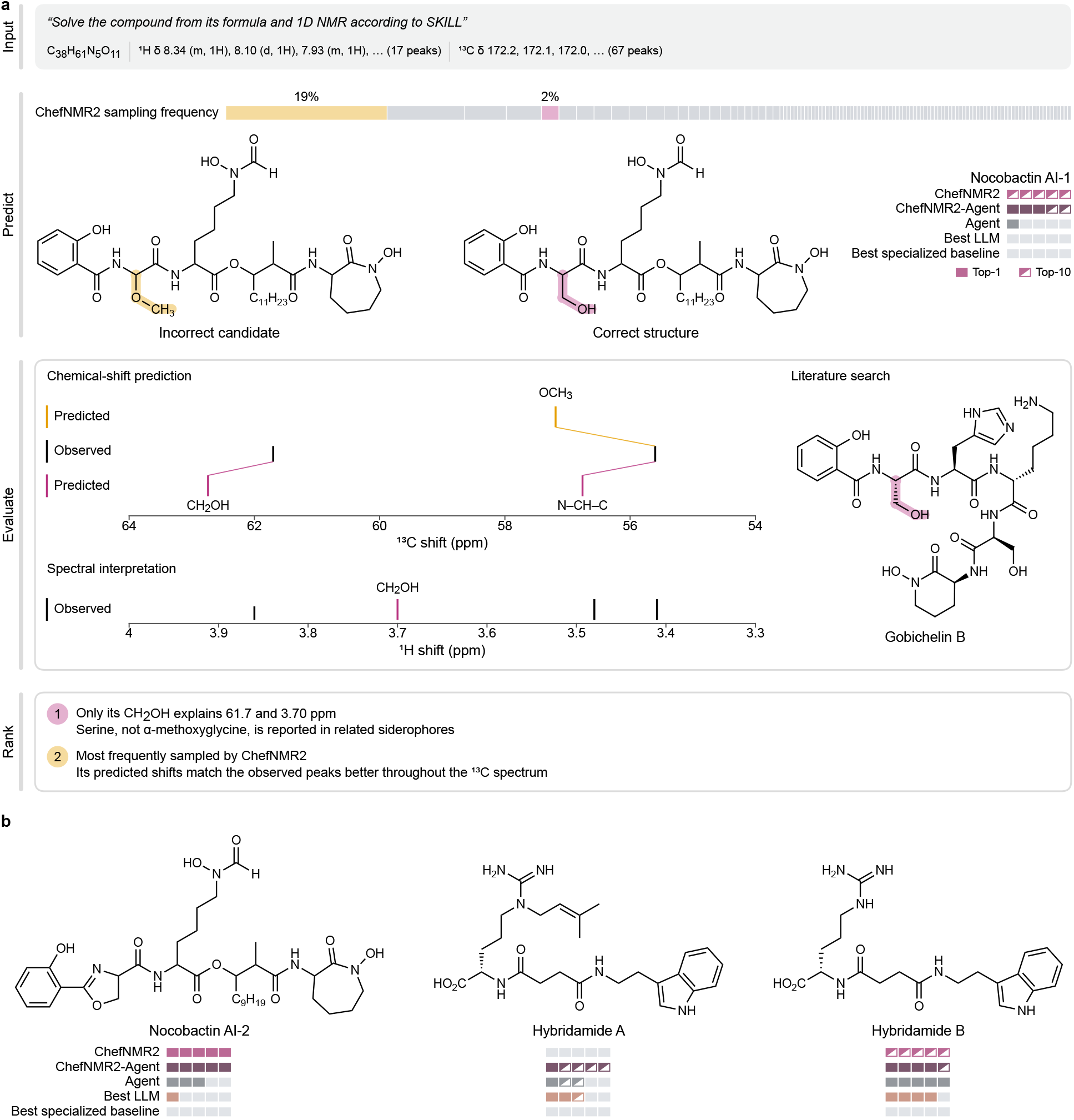
Elucidation of four novel natural products by ChefNMR2 - Agent. **a**, Example workflow for nocobactin AI-1. Input, molecular formula, ^1^H and ^13^C NMR peak lists and the skill (ChefNMR2 - Agent’s instructions). Predict, candidate structures generated by ChefNMR2, with bar widths showing their sampling frequencies. The incorrect candidate (amber) and correct structure (pink) are shown with their differing groups highlighted. In Evaluate, the agent compares predicted ^13^C shifts of the two structures (matching colors) with the observed peaks (middle), with lines joining matched peaks. Spectral interpretation marks the observed ^1^H peak that the agent assigns to the CH_2_OH of the correct structure (pink). Literature search shows a related siderophore that the agent cites, with the group it shares with the correct structure highlighted. The agent interprets the ^1^H peaks and searches the literature while ranking. Rank, ChefNMR2 - Agent’s top two candidates and summaries of the key reasons for their ranking. Each row of squares shows one system’s five runs, best first. Top-1, the correct structure ranked first. Top-10, ranked second to tenth. Open squares, not in the top ten or no answer. Structures are matched by planar structure. Agent, the same agent platform without the skill, ChefNMR2 or chemical-shift prediction tools. Best LLM, the most accurate of the six LLMs on these molecules (Claude-Opus-5). Best specialized baseline, the best rank among NMRTransformer, NMRPeak-G-Raw, UltraNMR and NMR-Solver in each run. **b**, Elucidation of nocobactin AI-2 and hybridamides A and B, shown as in **a**. Nocobactin AI-2 uses automatically picked peak lists without manual correction.

For the second analog, termed nocobactin AI-2, we use automatically picked peak lists without manual correction to test elucidation from noisy experimental inputs. ChefNMR2 ranks the correct structure first, but with a sampling frequency of 42%. After comparing the candidate with the spectra, ChefNMR2 - Agent retains it as the top-ranked structure in every run (Fig. 4b). In this case, the additional analysis supports a correct prediction despite uncertainty among the model’s samples.

We also tested two novel natural products, hybridamides A and B, that were recently isolated as part of a separate study from *Streptomyces* sp. HPH0547, the structures of which have not been reported ^52^. These were elucidated by human chemists using 1D/2D NMR data, and the 1D spectra were then provided to ChefNMR2. For hybridamide B, ChefNMR2 predicts the correct building blocks but inverts the order of succinate and arginine. It ranks this incorrect isomer first and the correct structure fourth to seventh. ChefNMR2 - Agent then obtains predictions from an independent LLM call, which predicts the correct form, and it switches the order of these two units and predicts the correct structure for hybridamide B.

We chose hybridamide A because it carries an unprecedented prenyl substitution at the *N^ε^*-nitrogen of arginine. Remarkably, ChefNMR2 captures all the components of hybridamide A accurately, but places the prenyl group on the succinyl amide nitrogen. By enumerating alternative prenyl group attachment sites and comparing succinyl carbonyl-carbon chemical shifts, the agent predicts the correct structure in every run and ranks it among its top two candidates in four of five runs, appropriately placing it on the guanidine *N^ε^*-nitrogen. These cases show how chemical reasoning and tool use can recover lower-ranked candidates and introduce structures missed by molecular generation. Moreover, they demonstrate the utility of this dual system in elucidating structures of natural products with novel chemotypes.

To assess time, accuracy, and cost involved in solving these four structures, we assembled data from human chemists in comparison with ChefNMR2 and ChefNMR2 - Agent. While the human chemist provides the correct structure using 2D NMR and tandem mass spectrometry data, ChefNMR2 is 1,032-fold faster with nocobactin AI-2. Meanwhile, the agent is approx. 86-fold faster for nocobactin AI-1, hybridamide A, and hybridamide B.

## 3. Discussion

In this work, we combine a state-of-the-art molecular generative model with emerging and complementary capabilities of scientific agents (Fig. 1b). The core generative model, ChefNMR2, is trained on an extensive corpus of molecular and spectral data and outperforms existing tools developed for NMR structure elucidation (Fig. 2). Recent advances in general-purpose LLMs have made it possible to approach the same task through natural language reasoning informed by broad chemical knowledge, including the ability to interpret NMR spectra ^24,25^. Accordingly, we perform a large-scale benchmark of frontier reasoning LLMs to assess their capabilities alongside specialized tools. ChefNMR2 achieves higher overall accuracy (Fig. 2c), better-calibrated confidence estimates (Fig. 2f), at up to 2,170-fold lower median runtime and 360-fold lower median cost per molecule than the LLMs (Fig. 2g–h).

We implement ChefNMR2 - Agent on the Kosmos agent platform ^30^, which coordinates chemistry and literature search subagents, with Claude-Opus-5.5 and GPT-6-Astra as base models (Methods 4.6.2). The agentic elucidation procedure is structured with a skill to guide molecular generation and evaluation (Fig. 1f), while allowing the agent to leverage its chemical reasoning capabilities. An agent given access to ChefNMR2 without the prescribed skill achieves comparable performance on the four unpublished compounds (Fig. S4), suggesting that useful elucidation strategies can also emerge without an explicitly defined workflow. We demonstrate the utility of ChefNMR2 - Agent by elucidating the structures of new natural products and an unusual substructure in hybridamide A. Even in cases where ChefNMR2 - Agent does not predict the correct structure, it significantly cuts the time needed to arrive at the final solution. Moreover, trained chemists could use additional data (such as tandem mass spectrometry or 2D NMR) to readily discern between top-ranked structures.

In its current form, ChefNMR2 uses the molecular formula as an auxiliary input, as it is typically the most readily obtainable from high-resolution mass spectrometry and effectively constrains the space of candidate structures, especially for large, complex molecules. Future models could relax this requirement by jointly generating atom counts, element identities and coordinates alongside conditioning on the spectra.

ChefNMR2 predicts planar structures without assigning stereochemistry. The 1D NMR inputs used to train the model do not generally provide sufficient information to distinguish stereoisomers, and chemists often rely on additional measurements, including 2D NMR correlations and chemical degradation analysis. Extending the model to use such information will require richer experimental datasets for retraining the generative model. These can potentially be obtained through a coordinated community effort. In parallel, ChefNMR2 - Agent offers a flexible interface for including additional compound-specific information. This flexibility accommodates the iterative, case-dependent nature of structure elucidation and allows agents to carry out structure elucidation tailored to specific scientific applications. Together, specialized generative models and scientific agents provide a foundation for collaboration between chemists and AI toward the grand challenge of automated structure elucidation.

## 4. Methods

### 4.1. Training data

ChefNMR2 is trained on three datasets that pair molecular structures with ^1^H and ^13^C NMR peak lists, one experimental and two simulated (Fig. 2a). NMRexp holds experimental peak lists that the NMRexp database ^36^ (version 2, file NMRexp_10to24_1_1004.parquet) extracted from the supporting information of published articles. SpectraNP+ holds peak lists that we simulate for natural products from COCONUT ^38^ (release of October 2024), NPAtlas ^39^ (release of September 2024) and NP-MRD ^40^ (downloaded in March 2025), greatly extending the SpectraNP set of our earlier work ^18^. USPTO holds the simulated peak lists that Alberts et al. ^21^ released for molecules from reactions in US patents ^37^. Sections 4.1.1 and 4.1.2 describe how structures and peak lists are processed, and Section 4.1.3 how the datasets are split.

#### 4.1.1. Structure processing

ChefNMR2 generates atomic coordinates and assigns bonds to them to obtain planar structures for ranking and evaluation. We compute up to ten conformers of each molecule in the datasets with the ETKDGv3 method ^53,54^ in RDKit ^55^ (version 2025.09.4), each optimized with the MMFF94 force field ^56^ or, if that fails, with UFF ^57^. The DetermineBonds function ^58^ of RDKit infers the bonds from the elements and coordinates. A conformer is kept only if its assigned bonds and stereochemistry match the recorded structure.

We retain molecules composed only of C, H, O, N, S, P, F, Cl, Br, I, B, Si, As and Se, with at most 200 atoms including hydrogens. Each retained molecule is neutral and single-component, with at least one kept conformer and a peak list for at least one nucleus.

#### 4.1.2. NMR peak-list processing

Published NMR data, such as those collected in the NMRexp database ^36^, are usually reported as peak lists, and the original free induction decays are rarely available ^59^. We therefore use peak lists, which also limit the effect of baseline distortion and of large differences in signal intensity in the raw spectra. We represent each ^1^H peak by its chemical shift, multiplicity label and number of protons, and each ^13^C peak by its chemical shift. We keep only peak lists whose shifts lie within −2 to 12 ppm for ^1^H and −20 to 230 ppm for ^13^C. To standardize multiplicity labels across sources, we write every label with the symbols s, d, t, q, p (quintet), h (sextet), hept (septet) and m, alone or combined, as in ddd or dt. We remove qualifiers such as apparent and broad, and write m for a label without a resolved splitting pattern, such as broad alone or overlapped. Simulated J coupling constants differ from those in experimental spectra, so we leave them out of every ^1^H peak list.

For NMRexp, we use the peak lists that the database ^36^ parsed from the text of each ^1^H and ^13^C record, and discard records whose shifts are not in monotonic order. We take the reported shift, or the midpoint of a reported range, as the shift of a ^1^H peak, and the reported integral as its number of protons. A ^1^H record is also discarded if a number of protons is not positive or a multiplicity label, such as ABX, AA^′^BB^′^ or nonet, cannot be standardized. We take each reported ^13^C shift as one peak.

Our SpectraNP+ peak lists are simulated for CDCl_3_ solvent with the Modgraph NMRPredict predictor in Mnova ^60^ (version 15.0.1). The ^1^H spectra are simulated at 500.13 MHz with a line width of 0.75 Hz. Each ^1^H peak list is the multiplet table of Mnova, from which we take the centroid of each multiplet as its shift, together with its multiplicity label and number of protons. The sum of Mnova’s integrals usually differs from the molecule’s hydrogen count. We discard the ^1^H lists of a molecule when this difference is large (above +20 or below −100 protons) or when a single peak has more protons than the whole molecule. The proton-decoupled ^13^C spectra are simulated at 125.03 MHz with a line width of 1.50 Hz. Each ^13^C peak list gives one shift per carbon atom, so symmetry-equivalent carbons appear as repeated shifts.

USPTO provides, for each molecule, the ^1^H and ^13^C peak lists that Alberts et al. ^21^ obtained with Mnova from simulated spectra. As for SpectraNP+, each ^1^H peak list is the multiplet table of Mnova, from which we take the centroid of each multiplet as its shift, together with its multiplicity label and number of protons. Each ^13^C peak list holds the peaks that Mnova picks from the simulated spectrum, so symmetry-equivalent carbons appear once and carbons coupled to fluorine or phosphorus appear as several peaks.

#### 4.1.3. Dataset splits

We split each dataset at random into training, validation and test sets in a ratio of about 85:5:10, or 90:5:5 for SpectraNP+. Most molecules with a peak list for only one nucleus are placed in the training set. Because ChefNMR2 elucidates planar structures, the split groups molecules by planar structure, so that stereoisomers fall in the same set. We identify each planar structure by its canonical SMILES string without stereochemistry ^61,62^. Because the three datasets share compounds, we also remove from each training set the molecules whose planar structures occur in the validation or test set of another dataset. The final training, validation and test sets hold 1,187,838, 56,082 and 123,801 molecules for NMRexp, 674,761, 34,226 and 35,129 for SpectraNP+, and 662,342, 34,730 and 78,645 for USPTO. Validation and test use only the molecules with both peak lists.

### 4.2. ChefNMR2 model

ChefNMR2 is a generative model for elucidating molecular structures from the molecular formula and the ^1^H and ^13^C peak lists (Fig. 1c). It generates atomic coordinates by flow matching (Section 4.2.1) with a network that consists of a peak encoder (Section 4.2.2) and a multimodal diffusion transformer (Section 4.2.3). Section 4.2.4 gives the training settings. Structures are ranked by sampling frequency, which also adapts the number of samples to each molecule to save computation (Section 4.2.5).

In ChefNMR2, we represent a molecule with *N* atoms by its three-dimensional coordinates **X** ∈ ℝ*^N^*^×3^ and the one-hot encoded element of each atom, **A** ∈ {0, 1}*^N^*^×^*^d^*^atom^ over *d*atom possible elements, taken from the molecular formula. The spectra S = (P_H_, P_C_) are two peak lists, in which P_H_ = {(*δ_i_*, *m_i_*, *n_i_*)} gives the chemical shift, multiplicity label and number of protons of each ^1^H peak, and P_C_ = {*δ_j_*} gives the chemical shift of each ^13^C peak. We formulate structure elucidation as sampling **X** from the conditional distribution *p*(**X** | **A**, S).

#### 4.2.1. Training and sampling of atomic flow-matching model

ChefNMR2 generates atomic coordinates by flow matching ^34,63^, which learns a velocity field whose flow transforms the Gaussian distribution into *p*(**X** | **A**, S). To learn this field, we construct a straight path **X***_t_* = (1 − *t*) **X**_0_ + *t* **X**_1_, *t* ∈ [0, 1], between a conformer **X**_0_ of the molecule and Gaussian noise **X**_1_ ∼ N (**0**, **I**), and train the network **v***_θ_*(**X***_t_*, *t*, **A**, S) to predict its velocity **X**_1_ − **X**_0_. The training loss is

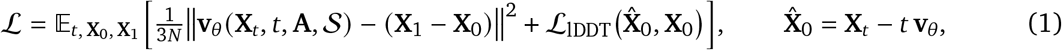

where *t* is drawn from a logit-normal distribution ^33^ shifted toward low noise and **X**^^^0 is the estimate of **X**_0_ implied by the predicted velocity. The term L_lDDT_ is a smooth local distance difference test (lDDT) loss adapted from AlphaFold 3^35,64^, which penalizes errors in the interatomic distances of **X**^^^0 (Algorithm S1). The training procedure and its settings are provided in Algorithm S2. Samples are generated by drawing **X**_1_ from N (**0**, **I**) and evolving it from *t* = 1 to *t* = 0 according to the stochastic differential equation ^65^

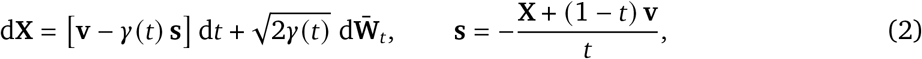

where **v** is the velocity, **s** is the score ∇**_X_** log *p_t_* (**X** | **A**, S) expressed in terms of **v**, **W̄***t* is a reverse-time Wiener process and *γ*(*t*) is a time-dependent diffusion coefficient.

Instead of using the learned velocity **v***_θ_*(**X**, *t*, **A**, S) directly as **v** in equation (2), we use classifier-free guidance ^66^, which is widely used to improve sample quality in diffusion and flow-based models ^65,67^. During training, both NMR spectra are randomly dropped, so that the network also learns to predict the velocity conditioned on the molecular formula alone, **v***_θ_*(**X**, *t*, **A**, ∅). During sampling, the two predictions are combined as

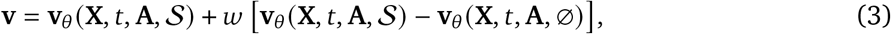

where *w* is the guidance scale. The sampling procedure is provided in Algorithm S3.

#### 4.2.2. Peak encoder

The network **v***_θ_* encodes the spectra S with a peak encoder, which represents each peak as one token. A ^1^H chemical shift *δ* is encoded by Fourier features ^68^ at *K* fixed, logarithmically spaced frequencies *f_k_*, which are sensitive to both large and small differences in shift, followed by a multilayer perceptron (MLP),

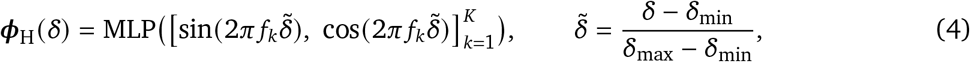

where *δ*_min_and *δ*max are the preset minimum and maximum ^1^H chemical shifts. The token of ^1^H peak *i* combines the embedding of its shift *δ_i_* with its multiplicity label *m_i_* and number of protons *n_i_*,

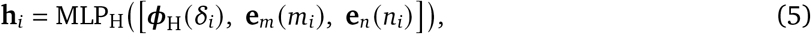

where **e***_m_* and **e***_n_* are learned embeddings and [·] denotes concatenation. Simulated ^1^H peak lists contain many detailed multiplicity labels that experimental lists rarely use, so ChefNMR2 keeps eight common multiplicity classes (s, d, t, q, dd, dt, td and m) and merges every other label into m. Numbers of protons are rounded to the nearest integer, and the few values above 20 are set to 20. The token of ^13^C peak *j* is the embedding of its shift alone, **c** *_j_* = ***φ***_C_ (*δ_j_*), computed as in equation (4) with its own chemical shift range, Fourier feature frequencies and MLP. A learned embedding of the nucleus is added to each token. Because the peaks of a spectrum form an unordered set, no positional encoding is used.

A stack of transformer blocks updates the ^1^H and ^13^C tokens together by self-attention, so that peaks of the two nuclei attend to each other. Attention pooling with a learned summary token ^69,70^ then aggregates the updated tokens into a single spectral embedding vector **z**_S_. Both the updated tokens and **z**_S_ condition the multimodal diffusion transformer. The encoder settings are provided in Table S2.

#### 4.2.3. Multimodal diffusion transformer

The network **v***_θ_* predicts the velocity with a multimodal diffusion transformer (MMDiT) ^33^. Each atom token is computed by an MLP from the atom’s noisy coordinates and one-hot element (its rows of **X***_t_* and **A**), and the peak tokens are the updated tokens of the peak encoder. In every block, atom and peak tokens have separate weights but share one self-attention operation, so every atom attends to every peak and vice versa. The sum of an embedding of the time *t* and the spectral embedding **z**_S_ conditions the atom and peak tokens of every block through adaptive layer normalization with zero initialization (adaLN-Zero) ^67^. A final adaLN-modulated linear layer maps each atom token to the velocity of that atom. The network is not equivariant to rotations of the molecule, so it is trained on conformers rotated at random to learn this symmetry (Algorithm S2), as in AlphaFold 3 and other non-equivariant generative models of molecules ^35,71,72^. The network settings are provided in Table S2.

#### 4.2.4. Training settings

We train ChefNMR2 jointly on the three datasets for 1,400 epochs, using the Adam optimizer ^73^ with a constant learning rate of 10^−4^ and a batch of 1,024 molecules. Throughout training, we keep an exponential moving average of the weights and use it as the trained model.

The three datasets differ in size and in whether their spectra are experimental or simulated. Each molecule of a batch is drawn at random from one dataset, chosen with equal probability, so each dataset supplies a third of the batch on average, whatever its size. One of its stored conformers and one of its peak lists per nucleus are then chosen at random, so that training covers the different conformers and peak lists of each molecule. To make ChefNMR2 tolerant of noise in experimental ^13^C peak lists, we perturb each simulated list with random peak shifts, extra peaks near existing ones and solvent peaks (Algorithm S4). The detailed training settings are provided in Table S2.

#### 4.2.5. Calibrated confidence and adaptive sampling

A well-calibrated generative model needs no separate confidence module, because the sampling frequency of a structure then estimates its probability of correctness. We therefore draw many samples for each molecule, convert each set of three-dimensional atomic coordinates into a planar structure by the bond assignment of Section 4.1.1, and rank the structures by this sampling frequency. ChefNMR2 predicts the top-ranked structure, whose sampling frequency is a well-calibrated confidence (Fig. 2f).

To avoid spending computation on molecules for which it is already confident, ChefNMR2 uses the sampling frequency to decide how many samples to draw. Sampling terminates early if a single structure accounts for nearly all of the structures obtained from the initial samples. Otherwise, additional samples are drawn to increase the chance of observing structures with a low sampling frequency. The number drawn is largest when nearly every structure obtained is distinct, or when none is obtained. All samples are drawn in equal numbers from the ten checkpoints saved during the last 500 epochs of training, so that no single checkpoint has to be chosen. The procedure and its settings are provided in Algorithm S5.

### 4.3. Benchmark curation

The benchmark comprises five test sets, held out from the training datasets or curated from public NMR databases (Section 4.3.1). It also includes natural products with published structures (Section 4.3.2) and natural products whose structures are not yet published (Section 4.3.3). Every structure is processed as in Section 4.1.1, and its peak lists are written in the form of Section 4.1.2.

#### 4.3.1. Held-out and public-database sets

Each held-out set holds 500 molecules with distinct planar structures drawn at random from the test set of NMRexp, SpectraNP+ or USPTO (Section 4.1.3).

To test whether a system can elucidate small compounds from automatically picked peak lists, we use SpecTeach, whose ^1^H and ^13^C free induction decays were recorded for undergraduate teaching ^44^. We keep the processing supplied with each free induction decay in the released Mnova files. Mnova’s automatic multiplet analysis (^1^H) and peak picking (^13^C) produce peak lists with common experimental noise, including solvent and impurity peaks, split or missed peaks, and inaccurate numbers of protons or multiplicity labels.

We read each ^1^H multiplet table as for SpectraNP+ (Section 4.1.2). We remove only compounds whose lists belong to another compound or whose ^1^H or ^13^C list contains none of the compound’s peaks. After processing, 216 compounds with both peak lists remain. NMRShiftDB2 tests elucidation from assigned peak lists in the public nmrshiftdb2 database ^42,43^ (nmrshiftdb2withsignals.sd, downloaded in September 2026). We select records with experimental ^1^H and ^13^C peak lists and a multiplicity for every ^1^H peak. The records give no integrals, so we use the number of hydrogens assigned to each ^1^H peak as its number of protons. We remove records whose peak lists are incomplete or do not match the structure. After processing, 315 compounds with distinct planar structures and both peak lists remain.

To compare the five sets, we compute for each set the distributions of heavy-atom count and of ^1^H and ^13^C shifts (Fig. 2b). Each shift distribution pools the peaks of all molecules in a set, weighted so that every molecule contributes equally. Each ^13^C peak of a molecule has the same weight, and each ^1^H peak a weight proportional to its number of protons. We use Gaussian kernel density estimates, with bandwidths chosen by leave-one-out cross-validation for shifts (2.0 ppm for ^13^C, 0.08 ppm for ^1^H) and by Scott’s rule for heavy-atom count.

#### 4.3.2. Well-known natural products and their derivatives

To test whether ChefNMR2 can elucidate complex natural products, we use nine well-known examples from diverse structural classes, with 20 to 80 heavy atoms (Fig. 3a,b). They are artemisinin ^74^, ginkgolide C ^75^, ingenol ^76^, lysergic acid ^77^, morphine ^78^, penicillin G, rapamycin ^79^, taxol ^80^ and the vancomycin aglycone ^81^. The spectra of penicillin G were recorded in D_2_O at the Princeton University Department of Chemistry NMR Core Facility (Bruker 500 MHz NMR spectrometer equipped with a DCH double resonance cryoprobe). We process the free induction decays in Mnova, pick the peaks and integrate the ^1^H spectrum manually, and take the multiplicity labels from its automatic multiplet analysis. The resulting tables are read as for SpecTeach (Section 4.3.1). For the other eight, we take the ^1^H and ^13^C peak lists, with multiplicities and integrals, from the supporting information of published articles.

To test whether ChefNMR2 can elucidate closely related structures, we use seven taxane diterpenoids from a study of taxol biosynthesis ^46^ (Fig. 3b,c). The supporting information of this study assigns their ^1^H and ^13^C shifts in CDCl_3_ atom by atom. We build their peak lists from these tables, merging carbons that share a shift and protons that share a shift and multiplicity, and using m where the table gives no multiplicity.

#### 4.3.3. Novel natural products

Nocobactin AI-1 and AI-2 were isolated and structurally elucidated in this study. Hybridamides A and B were available from a previous study ^52^. To obtain nocobactin AI-1 and AI-2, we began by culturing *N. transvalensis* NRRL B-16037 on Bennett’s agar. A single colony was then used to inoculate 3 mL of Bennett’s broth in a 14-mL culture tube. After growing for 3 days at 30 °C/250 r.p.m., 1 mL of the initial culture was used to inoculate 100 mL Bennett’s broth in each of two 250-mL Erlenmeyer flasks and cultured at 30 °C/250 r.p.m. for another 3 days. Cell pellets were then collected by centrifugation (30 min, 4,600*g*, 4 °C) and diluted into 8 × 1 L of R4 media in 2.8-L Fernbach flasks to a final concentration of 0.05% (wt/v), and cultured at 30 °C/220 r.p.m. for 10 days. The culture mixture was spun down (60 min, 18,000*g*, 4 °C), and the supernatant was loaded onto a prepacked C18 column (Phenomenex, 50 µm, 65 Å, 10 g). Elution was carried out with water and then 20%, 40%, 60%, 80%, and 100% MeCN in water stepwise. Analysis of the fractions by high performance liquid chromatography-coupled mass spectrometry (HPLC-MS) showed that the 80% and 100% fractions contain the two new nocobactins. These fractions were combined and dried *in vacuo*, resuspended in 50 mL 80% MeCN in water (+0.1% formic acid), and further purified by preparative HPLC with a Luna C18 column (Phenomenex, 5 µm, 21.2 × 250 mm) operating at a flow rate of 16 mL/min with mobile phases consisting of water and MeCN (+0.1% formic acid). After injection, elution was carried out isocratically with 50% MeCN in water for 2 min, followed by a gradient of 50%–95% MeCN in water over 22 min, and a gradient of 95%–100% MeCN over 10 min. Fractions were collected in 1-min intervals over the time range of 8–30 min. Fractions containing the target nocobactins, as judged by HPLC-MS analysis, were combined, dried *in vacuo*, resuspended in 80% MeCN in water and further purified on a semipreparative HPLC system. The sample was applied to a Luna phenyl-hexyl column (Phenomenex, 5 µm, 10.0 × 250 mm) operating at a flow rate of 2.5 mL/min with the same mobile phases as above and a gradient of 80–90% MeCN in water over 40 min, followed by a gradient of 90–100% MeCN over 10 min. Fractions containing pure nocobactins were combined and lyophilized to dryness. This procedure yielded 2.1 mg of nocobactin AI-1 and 4.5 mg of nocobactin AI-2. The NMR spectra of nocobactin AI-1 and AI-2 were recorded in DMSO-*d*6; those for hybridamides A and B were previously recorded in CD_3_OD.

### 4.4. Baselines and inference settings

We compare ChefNMR2 (Section 4.4.1) with specialized baselines for structure elucidation (Section 4.4.2) and general-purpose reasoning large language models (LLMs; Section 4.4.3). Each system receives the molecular formula and the NMR spectra, and returns a ranked list of candidate structures.

#### 4.4.1. ChefNMR2 inference settings

ChefNMR2 samples by the adaptive procedure of Section 4.2.5 and ranks the resulting structures by sampling frequency. On the five sets of Section 4.3.1, which hold 216 to 500 molecules each, we use one run of ChefNMR2 (Fig. 2). On the three sets of natural products (Sections 4.3.2 and 4.3.3), which hold 4 to 9 molecules each, we use five runs with different random seeds (Figs. 3 and 4).

#### 4.4.2. Specialized baselines

To compare ChefNMR2 with other specialized tools for structure elucidation, we use recent *de novo* and retrieval methods as baselines. The *de novo* baselines generate atomic coordinates (Chef NMR1^18^) or SMILES strings (NMRTransformer ^11^, NMRPeak ^13^ and UltraNMR ^15^). The retrieval baseline, NMR-Solver ^19^, searches a database of molecules with predicted shifts and refines the best matches by fragment-based optimization. We run each with its released model and default settings unless stated otherwise.

Chef NMR1 is our earlier atomic diffusion model, which reads NMR spectra sampled on a fixed chemical-shift grid rather than peak lists. It also conditions every atom on one pooled spectrum embedding instead of on individual peaks, and is trained as a diffusion model ^82^, whereas ChefNMR2 uses flow matching. It has 462 million parameters, compared with 288 million for ChefNMR2. We evaluate Chef NMR1 only on the test sets that provide full spectra, not just peak lists. We draw ten samples per molecule in a single run and report the structures in sampling order.

NMRTransformer is a multimodal transformer that generates SMILES strings from the molecular formula and any combination of ^1^H NMR, ^13^C NMR and infrared spectra. Its released checkpoint is pretrained on simulated USPTO spectra ^21^. We give it the molecular formula, padded to 15 tokens as in its training, and the NMR peak lists. As the model expects a range for each ^1^H shift, we set its upper and lower bounds to the shift centroid. It generates ten candidates by deterministic beam search.

NMRPeak-G-Raw, the generation module of NMRPeak, is a sequence-to-sequence transformer that generates SMILES strings from the molecular formula and ^1^H and ^13^C peak lists. It is trained on experimental peak lists from the NMRexp database that include J coupling constants. During inference, we give it the ^1^H peaks without J coupling constants. It generates ten candidates by deterministic beam search.

UltraNMR is a transformer that generates SMILES strings from the molecular formula and the ^1^H and ^13^C shifts. It is pretrained on spectra simulated for PubChem molecules, by NMRNet and by density functional theory, and its released *de novo* checkpoint is fine-tuned on experimental spectra of the NMRGym benchmark. It reads only the shift of each peak, not its multiplicity or number of protons. It generates ten candidates by deterministic beam search.

NMR-Solver is a retrieval method that refines its best database matches by fragment-based optimization. The observed ^1^H and ^13^C shifts are compared with a database of spectra simulated by NMRNet ^83^, a neural network trained on nmrshiftdb2, for about 106 million PubChem molecules. Both retrieved and refined structures are ranked by the agreement between their predicted shifts and the observed shifts. The highest-ranked candidates with the exact formula, up to ten, are reported.

#### 4.4.3. Reasoning large language models

To test whether general-purpose LLMs can elucidate structures from NMR data through their own chemical knowledge and reasoning, without tools or web access, we evaluate Claude-Opus-5, GPT-5.6-Sol, Gemini-3.6-Flash, GLM-5.2, Kimi-K3 and DeepSeek-V4-Flash. All models receive the same prompt, which gives the molecular formula and the NMR peak lists (Table S3). Each is asked to return a ranked list of up to ten SMILES strings, each with a confidence between 0 and 1. All models use the reasoning settings in Table S3, a one-hour timeout per request and their maximum output length, capped at 128,000 tokens. We query each model once per molecule because of the high cost. On the four novel natural products, we query each model five times (Fig. 4). The models were queried between 29 July and 26 October 2026.

### 4.5. Evaluation

To compare systems that return structures in different forms, we score all of them by one matching accuracy (Section 4.5.1). We then assess what each training set contributes to ChefNMR2 (Section 4.5.2). We also check whether a system may have seen a benchmark molecule in training (Section 4.5.3), and relate accuracy to similarity, confidence, and runtime and cost (Sections 4.5.4– 4.5.6).

#### 4.5.1. Matching accuracy

We compare planar structures, since stereochemistry cannot be reliably assigned from 1D NMR spectra without coupling constants or nuclear Overhauser effect correlations. Top-*k* matching accuracy (*k* = 1 or 10) checks whether the correct structure is exactly matched by any of the top-*k* candidates of a system (Section 4.4), as RDKit canonical SMILES strings without stereochemistry. A system that returns no usable structure counts as incorrect, including invalid generated coordinates or SMILES strings and LLM answers that are missing or reach the output limit.

#### 4.5.2. Contribution of each training set

We characterize the chemical diversity and structural complexity of the three training sets by the proportions of molecules containing different elements and ring structures (Fig. S1a,b). We classify ring structures by size, aromaticity, heteroatoms and how the rings are connected. Rings are identified using the symmetrized smallest set of smallest rings in RDKit ^55^.

To assess the benefit of combining training sets, we compare ChefNMR2 with models of the same architecture trained on each set alone (Fig. S1c). All models use the same training and inference settings and the same number of training batches.

We further test whether combining training sets makes ChefNMR2 more robust to experimental noise. We compare the four models’ top-1 accuracy on SpecTeach molecules grouped by the type of noise in their peak lists (Fig. S1d). The four groups have noise in the ^1^H peak list only, the ^13^C peak list only, both or neither. A ^13^C list counts as noisy if it contains more peaks than chemically distinct carbons, with diastereotopic carbons counted separately. A ^1^H list counts as noisy if its numbers of protons sum to fewer than the molecule’s C–H protons or more than its total hydrogens. The C–H count excludes hydrogens bonded to O, N or S, since experimental lists may omit exchangeable protons.

We also test whether combining training sets improves generalization to dissimilar molecules. For each NMRShiftDB2 molecule, we calculate its maximum Tanimoto similarity ^84^ to each training set using Morgan fingerprints ^85^ of radius 2 and 2,048 bits, without stereochemistry. Molecules are grouped by which training set gives the highest similarity, with ties included in each group. Molecules with similarity below 0.6 to all three training sets are considered dissimilar to the training data and grouped separately. We compare the four models’ top-1 accuracy within each group (Fig. S1f).

#### 4.5.3. Molecules seen in training

Accuracy on molecules seen in training can reflect memorization rather than elucidation. A benchmark molecule counts as seen in training if a system was trained on its planar structure with spectra of the same kind: both experimental, or both simulated by the same simulator. No benchmark molecule is seen in training by ChefNMR2. Four occur in its training data with simulated spectra, the taxanes taxusin and 15-hydroxy-11(15→1)-abeo-taxusin (Section 4.3.2), and taxol and penicillin G, written as tautomers in which the amide group is an imidic acid.

We mark a specialized baseline on a test set when at least 5% of the set is seen in its training data (Fig. 2c). For the *de novo* elucidation baselines, we use the training sets of the released models. The retrieval baseline, NMR-Solver, ranks its candidates with shifts predicted by NMRNet (Section 4.4.2). We therefore treat the experimental shifts used to train NMRNet as the training data of NMR-Solver. We compare the top-1 accuracy of each marked baseline on its seen and unseen molecules (Fig. S2a).

The accuracy of the LLMs can also reflect recall of published structures rather than elucidation. Their training data are enormous and undisclosed, so we look for recall in their answers instead of checking the overlap. We compare the answers of Claude-Opus-5, the most accurate LLM on the five test sets, on taxol and on two less well-known taxanes, and give further answers that name the molecule (Table S1).

#### 4.5.4. Similarity to training data

To test how well ChefNMR2 generalizes to molecules unlike those it was trained on, we compute the largest Tanimoto similarity of each benchmark molecule to any training molecule. The Tanimoto similarity of two molecules is the number of bits set in both of their fingerprints divided by the number set in at least one ^84^. We use Morgan fingerprints ^85^ of radius 2 and 2,048 bits, without stereochemistry. For each specialized baseline, we compute the same similarity to its own training data (Section 4.5.3), adding the PubChem molecules of the NMR-Solver database for UltraNMR and NMR-Solver. We compare how the top-1 accuracy of each baseline and of ChefNMR2 changes with this similarity (Fig. S2b).

#### 4.5.5. Calibration

We assess the calibration of the confidence reported by each system: the sampling frequency of the top-ranked structure for ChefNMR2 (Section 4.2.5) and Chef NMR1, the probability of the top-ranked structure for chemical language models (NMRPeak-G-Raw, NMRTransformer and UltraNMR), and the confidence that each LLM assigns to its first candidate. For chemical language models, we multiply the probabilities of the predicted tokens of each candidate at temperature 1 and add the products for candidates that represent the same structure. For each system, we divide the molecules it answers in the five test sets into six confidence bins, (0, 0.2], (0.2, 0.4], (0.4, 0.6], (0.6, 0.8], (0.8, 0.95] and (0.95, 1], and compare the mean confidence in each bin with the top-1 accuracy in that bin (Fig. 2f). For NMR-Solver, whose ranking score is the agreement between predicted and observed shifts rather than a probability, we show accuracy against this score only (Fig. S2f). The expected calibration error (ECE) ^45^ is the mean absolute difference between these two quantities, weighted by the number of molecules in each bin. Figs. 2f and S2f omit bins with fewer than ten molecules, which still count in the ECE. We apply the same analysis to the candidates ranked second and lower by each system except NMR-Solver, dividing the bin (0, 0.2] further at 0.01, 0.02, 0.05 and 0.1 because most sampling frequencies of the structures that ChefNMR2 ranks second and lower lie below 0.2 (Fig. S2f).

#### 4.5.6. Runtime and cost

We report the per-molecule runtime and cost of each system as medians with interquartile ranges (Fig. 2g,h). ChefNMR2 and the specialized baselines process molecules one at a time on NVIDIA H100 GPUs. Their runtime is the inference time excluding checkpoint loading, and their cost is estimated from the GPU time at Modal’s on-demand rate of US$3.95 per GPU-hour ^86^. Each specialized baseline runs on a single GPU. Each of the ten checkpoints of ChefNMR2 (Section 4.2.5) runs on its own GPU. Its runtime is that of the slowest checkpoint, as for a parallel run, and its GPU time is summed over all ten.

For the LLMs, runtime is the duration of the request that returned the response, and cost is estimated from the input and output tokens of that request, reasoning tokens included, at the list prices of Table S3.

### 4.6. ChefNMR2 - Agent

We develop ChefNMR2 - Agent to combine ChefNMR2’s predictions with an LLM agent’s chemical reasoning and use of other tools. We provide it as a skill, a set of instructions and scripts that the agent follows to predict, evaluate and rank candidate structures (Section 4.6.1). We evaluate ChefNMR2 - Agent on the four novel natural products of Section 4.3.3 using the Kosmos agent platform ^30^ (Section 4.6.2).

#### 4.6.1. Workflow and tools

The agent receives a molecular formula and ^1^H and ^13^C peak lists as text and is instructed to elucidate the compound’s planar structure following the skill. It parses the peak lists into a table of chemical shifts, multiplicity labels and numbers of protons. A script validates the input by checking the ^13^C peak count against the number of carbon atoms in the formula and the summed numbers of protons against the number of hydrogen atoms. It lists possible causes of any discrepancy, such as peak overlap or impurities, for the agent to further assess. The skill then guides the agent to predict, evaluate and rank candidate structures (Fig. 1f).

The agent collects candidate structures from ChefNMR2, an LLM, PubChem and enumeration of structural variants. It first calls ChefNMR2 using the checkpoints, settings and adaptive sampling procedure of Section 4.2.5 and Algorithm S5. If the initial samples meet the convergence criterion, it returns the most frequently sampled candidate without further evaluation. Otherwise, it expands sampling and adds candidates from the other sources. The LLM receives only the molecular formula, peak lists and the validation result, without the candidates generated by ChefNMR2. A script runs a database search in PubChem ^87^ for the molecular formula and keeps up to 200 compounds with their positions in the retrieved list. The agent also enumerates variants for each structural question that the spectra leave open, such as how fragments are joined, the site of a substituent or the size of a ring. It writes these questions from the spectra alone and fixes them before reading the candidates of the other sources. It then adds only the questions needed to include the first LLM proposal and every ChefNMR2 candidate with a sampling frequency of at least 5%. Scripts filter candidates for validity, neutral charge, a single fragment and agreement with the molecular formula, then merge duplicate planar structures and tautomers while preserving their sources.

The merged candidates are evaluated using spectral consistency tests and predicted chemical shifts. The agent writes a checklist of tests, such as whether the number of methyl groups is compatible with ^1^H peaks of three or more protons. A script counts the tests passed by each candidate. Each candidate is also evaluated with CSP5^41^, a model that predicts NMR chemical shifts from molecular structure. We compare its predicted ^13^C shifts with the observed peak list using a modified version of NMRPeak’s ^13^C spectrum similarity ^13^. Excess observed peaks are left unmatched and incur no penalty. A script answers each enumeration question by comparing the CSP5 scores of variants that differ only in that question, and leaves it undecided when these comparisons do not favor one answer.

The agent ranks candidates by reasoning about the spectra and candidate structures, including ChefNMR2’s sampling frequency, agreement between candidate sources, checklist results, CSP5 scores and the answers to the enumeration questions as evidence. The skill explains what this evidence measures and gives priority to diagnostic ^1^H and ^13^C evidence. The skill requires the returned list to include a candidate for every answer that the CSP5 comparisons leave undecided. When the list cannot hold them all, the agent covers questions about the skeleton first, such as how fragments are joined. The agent can request a literature search for the compound class or a structural motif when it could distinguish the top candidates, and use retrieved evidence to update its ranking. It returns up to ten ranked candidates, each with a written rationale.

#### 4.6.2. Agent platform and runs

We run ChefNMR2 - Agent on the Kosmos agent platform. An orchestrating agent delegates the workflow to subagents. One chemistry subagent performs prediction, evaluation and provisional ranking. When literature is needed, a literature subagent investigates focused questions and a second chemistry subagent updates the ranking. The orchestrating agent and the chemistry subagents run Claude-Opus-5.5. When Claude-Opus-5.5 refuses a request, the platform can send it to Claude-Opus-5 instead. The LLM proposals come from a separate API call to GPT-6-Astra at high reasoning effort. When GPT-6-Astra returns no proposal, the agent may ask Claude-Opus-5 instead.

We perform five runs per compound with ChefNMR2 - Agent and five with each of two controls (Figs. 4 and S4). Each run starts in a new session. To test how much the skill and its NMR tools improve elucidation, one control (Agent) runs the same agent platform without the skill, ChefNMR2 or CSP5. To separate the contribution of ChefNMR2 from that of CSP5 and the skill, the other control (ChefNMR2 - Agent, no skill) can call ChefNMR2 through the platform but works without the skill or CSP5. Both controls use the same platform and model and receive the same molecular formula and peak lists (Table S4). They are asked to return ten candidate planar structures and retain the platform’s other capabilities, including code execution and its built-in scientific subagents.

For each run, we report the rank of the correct structure in the returned list. The runs of ChefNMR2 - Agent took place between 10 and 12 October 2026, and the control runs on 19 October 2026.

To show how far the top-ranked candidate of each run lies from the correct structure, we call it correct, close or distant (Fig. S4). A candidate is close when it has the molecular formula of the correct structure and can be turned into it by breaking and forming at most four bonds, for example by moving a group or by turning an alcohol into an ether. Every other candidate is distant, as is a run without an answer. We also record the best class among the ten top-ranked candidates of each run. For the specialized baselines, we show the best rank among the four in each run (Figs. 4 and S4).

### 4.7. Chemical landscape analysis

To examine what ChefNMR2 learns, we extract its internal representations and compare them with the chemical space of the training data (Sections 4.7.1–4.7.3).

#### 4.7.1. Molecules and descriptors

We analyze 2,524,941 molecules from the three training datasets (Section 4.1). Each molecule is represented by 35 molecular descriptors computed with RDKit ^55^ and 37 spectral features. The spectral features summarize the peak lists that ChefNMR2 receives as input, such as the number of peaks and the range of chemical shifts for each nucleus.

#### 4.7.2. Embeddings of ChefNMR2

All embeddings are taken from the last of the ten checkpoints described in Section 4.2.5. The spectral embedding **z**_S_ produced by the peak encoder (Section 4.2.2) depends only on the peak lists and therefore reflects how the network represents NMR spectra. We compute **z**_S_ for every training molecule and for 16 reference molecules, comprising the natural products and taxanes (Section 4.3.2).

#### 4.7.3. Projections and principal components

Before projection, each dimension of the embedding is standardized to zero mean and unit variance, and the embedding is reduced to its first 50 principal components by principal component analysis (PCA). Reduced embeddings are mapped to two dimensions by uniform manifold approximation and projection (UMAP) ^88^ with 100 nearest neighbors, a cosine distance metric and initialization by PCA (random seed 0). The other settings are the UMAP defaults, including a minimum distance of 0.1 and a spread of 1.0. The projection of **z**_S_ is fitted jointly to the training and reference molecules (Figs. 2a and 3b).

To identify the main axes of variation in **z**_S_, we fit a separate eight-component PCA to the standardized embedding of the 2,524,941 training molecules and correlate each component (Pearson’s *r*) with the 35 RDKit descriptors and the 37 spectral features. Each of the first three components is plotted against the descriptor it correlates with most strongly, which is the number of heavy atoms (PC1), the fraction of sp^3^ carbons (PC2) and the number of aromatic carbocycles (PC3). Each is also plotted against one of three spectral features: the total number of peaks (PC1), the mean ^1^H (PC2), and ^13^C (PC3) chemical shifts. We take eight equally spaced values along the first two principal components from their 1st to 99th percentiles with the other component held at their median. For each value, we find the molecule closest to that point in the plane of the two components. Four of these molecules are shown with their structures and peak lists (Fig. S3).

## 5. Data availability

NMRexp (version 2) is available from https://doi.org/10.5281/zenodo.17296666. COCONUT (release of October 2024) is available from https://doi.org/10.5281/zenodo.13897048. NPAtlas (release of September 2024) is available from https://doi.org/10.5281/zenodo.13756408. NP-MRD is freely available at https://np-mrd.org/downloads. The simulated USPTO spectra of Alberts et al. are available from https://doi.org/10.5281/zenodo.11611178. The SpecTeach spectra are available from Van Bramer and Bastin ^44^. nmrshiftdb2 is available at https://nmrshiftdb.nmr.uni-koeln.de. The test sets as used in this study will be deposited on Zenodo, and SpectraNP+ will be released upon publication.

## 6. Code availability

The code of ChefNMR2 is available at https://github.com/ml-struct-bio/chefnmr2. Chef NMR1 is available at https://github.com/ml-struct-bio/chefnmr, with its checkpoints at https://doi.org/10.5281/zenodo.17766755. NMRTransformer is available at https://github.com/rxn4chemistry/MultimodalAnalytical, with its checkpoint at https://doi.org/10.5281/zenodo.16914550. NMRPeak is available at https://github.com/Colin-Jay/ NMRPeak, with its data and weights at https://doi.org/10.5281/zenodo.19122815. Ultra-NMR is available at https://github.com/wuycM/UltraNMR, with its checkpoints at https://huggingface.co/milesyc/ultranmr. NMR-Solver is available at https://github.com/YongqiJin/NMR-Solver, with its model and data at https://doi.org/10.5281/zenodo.16952024.

## 7. Acknowledgements

The authors thank Elizabeth Sattely, Franklin Wang, Ullrik Unneberg, Michael Albergo, and other members of the Zhong lab for helpful discussions. This work was supported by the Eric and Wendy Schmidt Transformative Technology Fund (E.D.Z. and M.R.S.), the Princeton Catalysis Initiative (E.D.Z. and M.R.S.), a Princeton SEAS Innovation grant (E.D.Z. and M.R.S.), the National Institutes of Health (DP2GM164606 to E.D.Z), Anthropic’s AI for Science Program (E.D.Z), the MacArthur Fellowship (M.R.S.), the Life Sciences Research Foundation (Postdoctoral Fellowship sponsored by the Open Philanthropy project to J.Z.H.), and the National Research Foundation of Korea (Postdoctoral Fellowship RS-2024-00406578 to J.S.A.). The authors acknowledge the use of computing resources at Princeton Research Computing, a consortium of groups led by the Princeton Institute for Computational Science and Engineering (PICSciE) and Office of Information Technology’s Research Computing.

## 8. Author contributions

E.D.Z. and M.R.S. conceived the project. Z.X. and E.D.Z. developed the method. Z.X. curated data, implemented the software, and performed the experiments, with contributions from Z.K. on LLM benchmarking, J.D.C. on chemical landscape analysis and baselines, and K.M. on code. Z.X., E.D.Z., Z.K., and A.D.W. developed ChefNMR2 - Agent. V.Y.Y., J.Z.H., and J.S.A. collected and elucidated natural product data and structures. Z.X., J.D.C., and R.C.H. prepared visualizations. Z.X., M.R.S., and E.D.Z. wrote the manuscript with input from all authors. E.D.Z., M.R.S., A.D.W., and S.G.R. supervised the project.

---

**Fig. S1.**
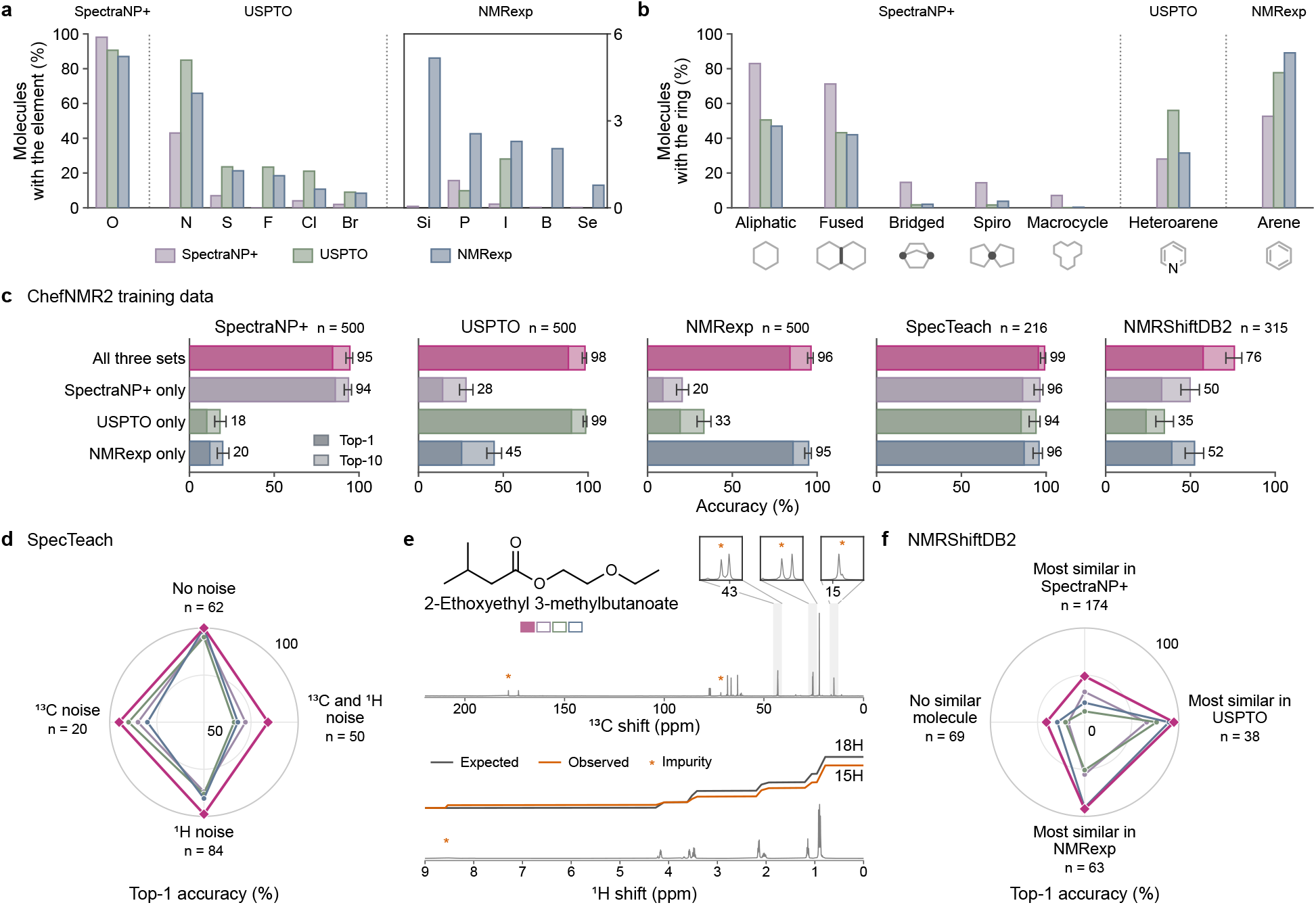
Contribution of each training set to accuracy and robustness to experimental noise. **a**, Share of each training set’s molecules containing each element besides C, H and As. Elements are grouped by the training set with the largest share. Bars inside the box are read on the right axis (0–6%). **b**, Share of molecules with each type of ring, grouped as in **a**. Macrocycle, 12 or more ring atoms. **c**, Top-1 (dark) and top-10 (light) accuracy of ChefNMR2 and of the same model trained on one training set alone. Numbers, top-10 accuracy. Error bars, 95% Wilson intervals of top-10 accuracy. **d**, Top-1 accuracy on SpecTeach by noise in its peak lists. ^13^C noise, more peaks than chemically distinct carbons. ^1^H noise, numbers of protons that sum to fewer than the compound’s C–H protons or more than its hydrogens. **e**, A SpecTeach example with experimental noise in both peak lists. Squares, the models of **c** in row order. Filled, correct structure ranked first. Open, not in the top ten. Asterisks, impurity peaks. Expected, the structure’s protons assigned to each peak. Observed, the numbers of protons in the automatically picked peak list. Both are added up from high to low shift. **f**, Top-1 accuracy on NMRShiftDB2 by the training set holding the most similar molecule (Tanimoto similarity ≥0.6). No similar molecule, below 0.6 in every training set. Ties are included in each group. Colors in **d**–**f** as in **c**.

**Fig. S2.**
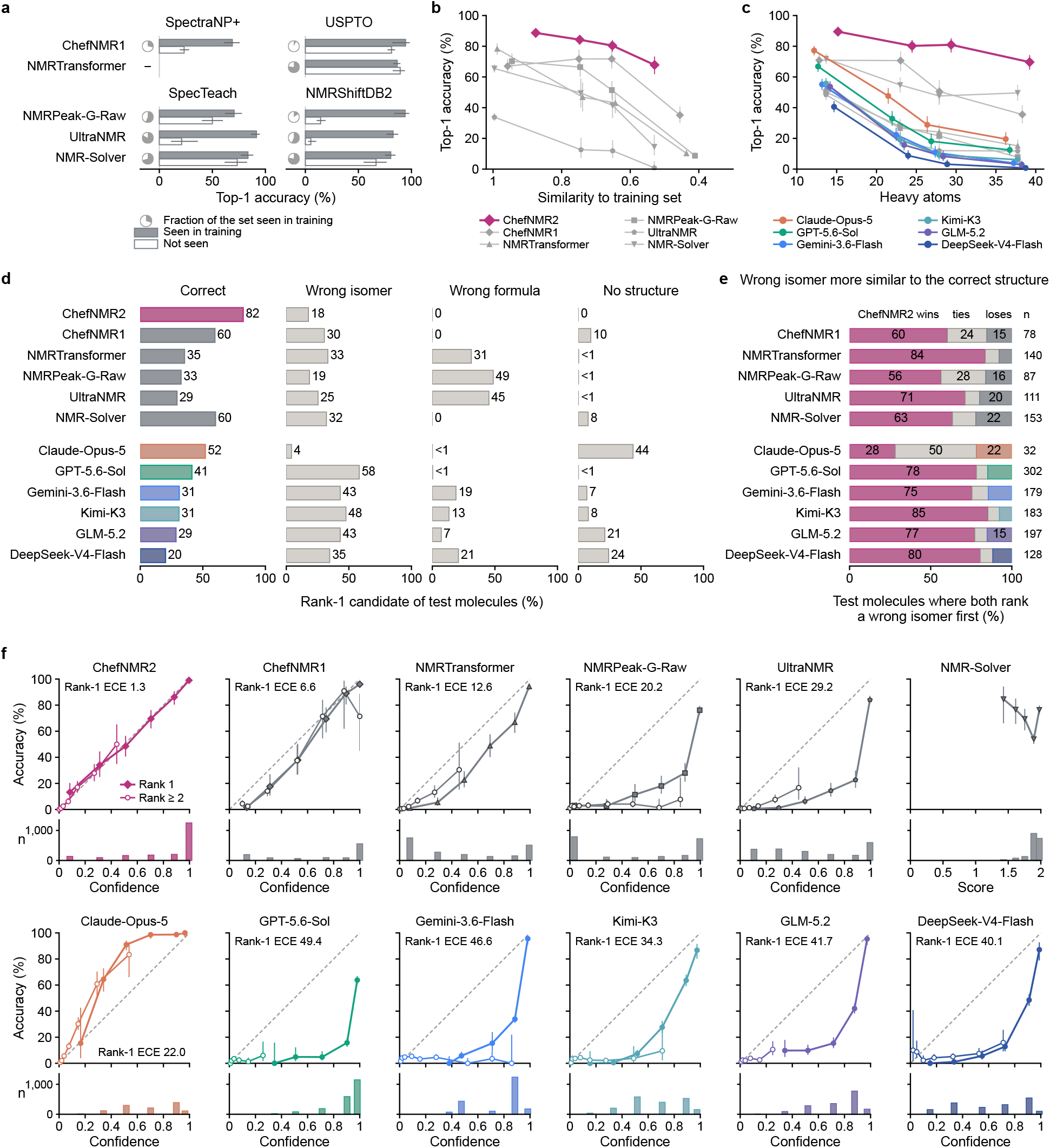
Generalization, failure cases and calibration across systems. **a**, Top-1 accuracy of each specialized baseline on test molecules of each set, seen in training (filled) or not (open). Pie, fraction of the set seen in training. –, less than 5% of the set seen in training. **b**,**c**, Top-1 accuracy by Tanimoto similarity to the nearest training molecule of each system (**b**) and by number of heavy atoms (**c**), each point at the mean of its interval. **d**, Class of the rank-1 candidate of each system. Wrong isomer, the molecular formula of the correct structure but another planar structure. No structure, no candidate or an invalid one. **e**, Test molecules on which ChefNMR2 and another system both rank a wrong isomer first. ChefNMR2 wins, ties or loses as its wrong isomer is more, equally or less similar to the correct structure (Tanimoto). *n*, number of test molecules. **f**, Accuracy against confidence of rank-1 (filled) and lower-ranked (open) candidates, over the molecules each system answers. Confidence is the sampling frequency (ChefNMR2, Chef NMR1), probability (chemical language models) or stated confidence (LLMs). For NMR-Solver, the agreement between predicted and observed shifts. Dashed line, confidence equal to accuracy. ECE, rank-1 expected calibration error. Bars, rank-1 candidates per bin. Test sets are pooled in **b**–**f**. Error bars, 95% Wilson intervals. In **d**, all 95% intervals are within 2.8 percentage points.

**Fig. S3.**
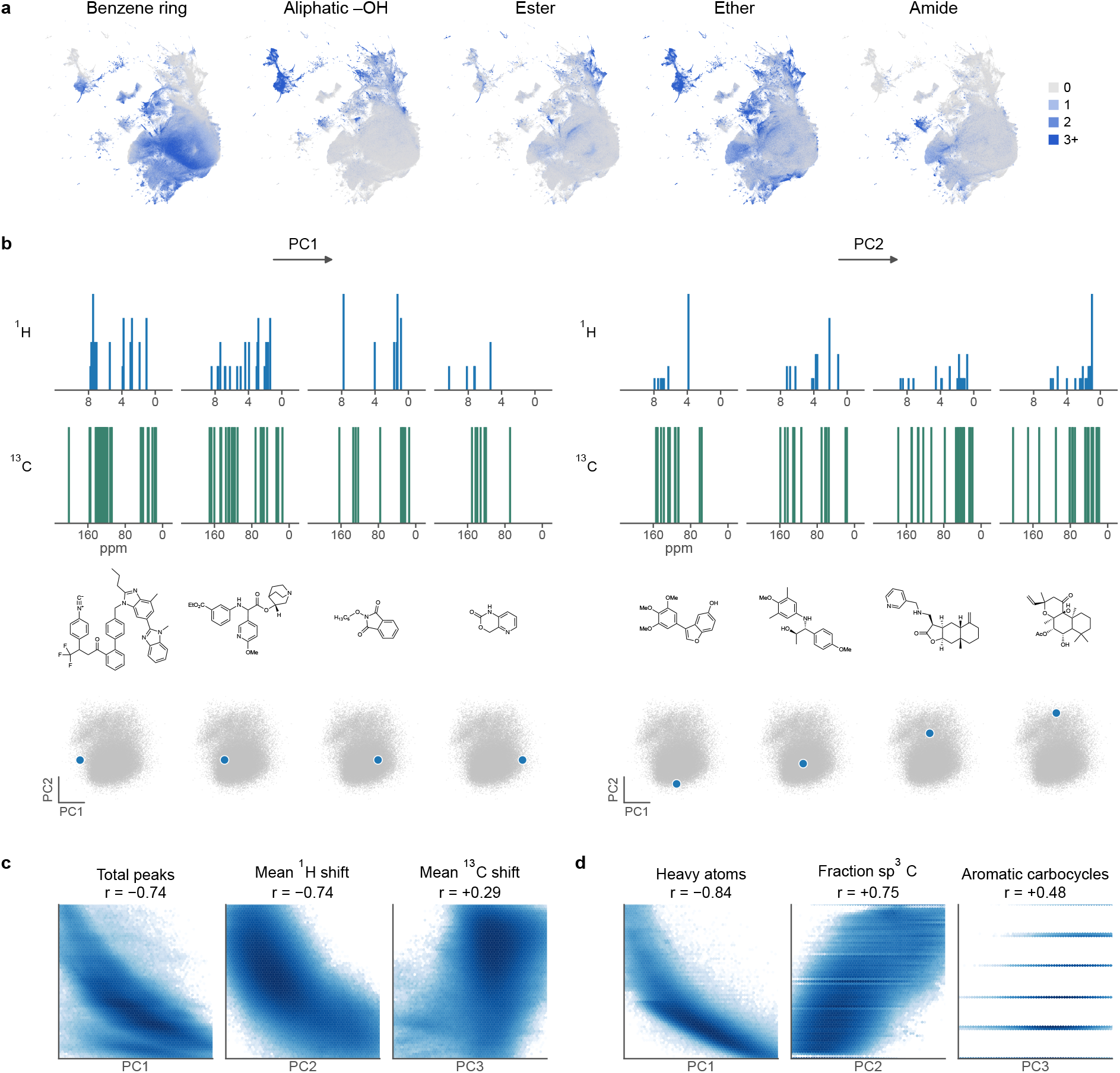
| **Correlating spectral embeddings with molecular descriptors. a**, UMAP of the spectral embeddings of the training molecules, colored by the number of each functional group. **b**, Traversals of PC1 and PC2, four of the eight steps of Methods 4.7.3. Steps span the 1st to 99th percentile of the axis. Spectra, structure and position in embedding space are shown at each step, for the nearest molecule with both a ^1^H and a ^13^C spectrum. **c**, Scatter plots of PC1 against the total number of peaks, PC2 against the mean ^1^H shift and PC3 against the mean ^13^C shift. **d**, Scatter plot of each PC against the molecular descriptor it correlates with most strongly. PC, principal component. *r*, Pearson correlation coefficient.

**Fig. S4.**
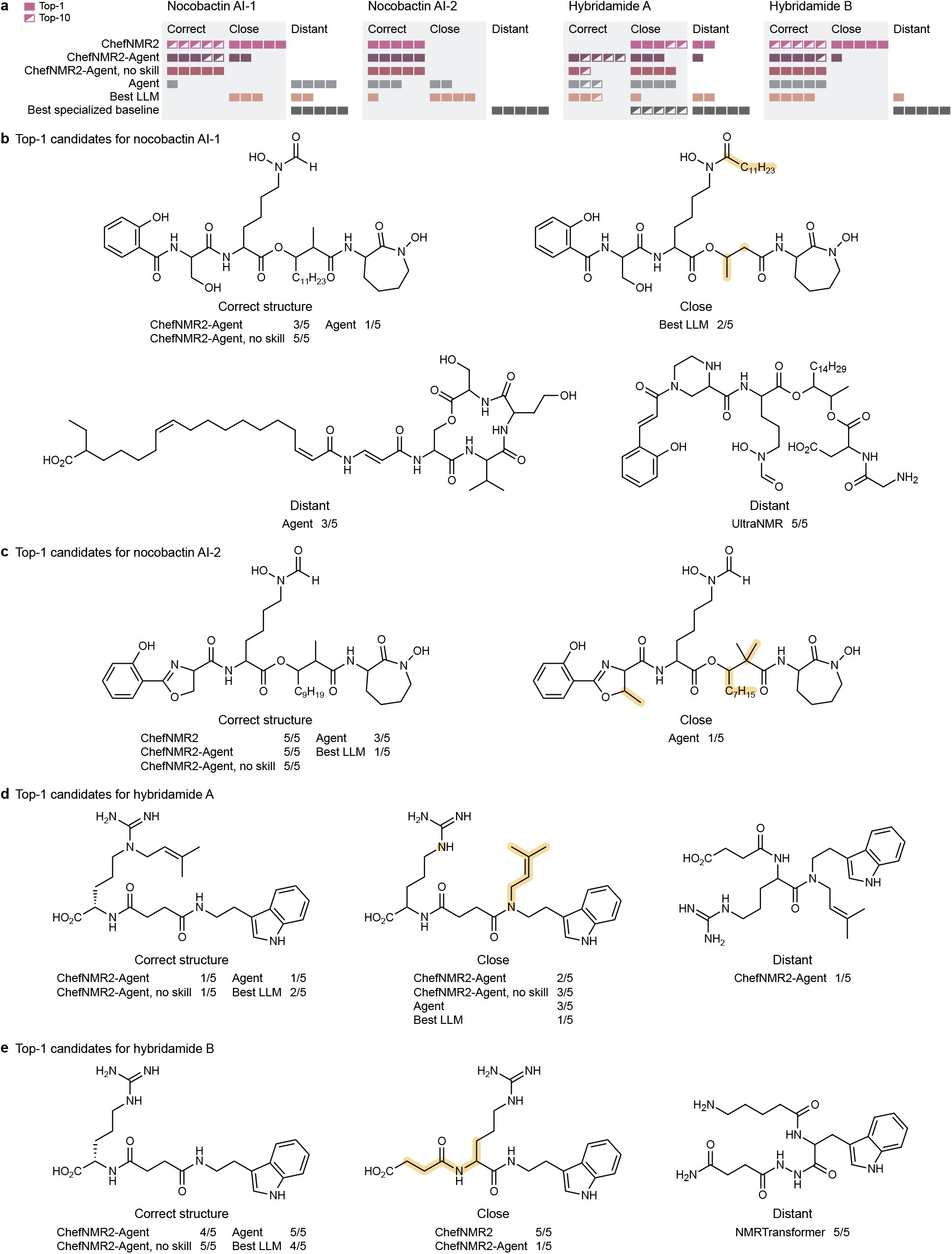
| **Top-1 candidates of each system for the four novel natural products. a**, Class of each run’s top-1 candidate. Close, a candidate with the molecular formula of the correct structure that can be turned into it by breaking and forming at most four bonds, for example by moving a group or by turning an alcohol into an ether. Distant, any other candidate, or no answer. Each system has five runs, and each square represents one run. Filled squares show the class of the top-1 candidate, and half-filled squares the best class among the top ten when it is better. ‘ChefNMR2 - Agent, no skill’ is the same agent platform with ChefNMR2 but without the skill (ChefNMR2 - Agent’s instructions) or chemical-shift prediction tools. Agent, the same agent platform without the skill, ChefNMR2 or chemical-shift prediction tools. Best LLM, the most accurate of the six LLMs on these molecules (Claude-Opus-5). Best specialized baseline, the best rank among NMRTransformer, NMRPeak-G-Raw, UltraNMR and NMR-Solver in each run. **b**–**e**, Selected top-1 candidates for nocobactin AI-1 (**b**), nocobactin AI-2 (**c**), hybridamide A (**d**) and hybridamide B (**e**). Under each structure are its class and the systems that rank it first, with the number of runs out of five. Highlights mark bonds that differ from the correct structure drawn in the same layout, and dots mark atoms that lost a substituent. Structures are matched by planar structure.

**Table S1.** LLM answers on well-known natural products and taxanes. a. , Answers that name the molecule, on the nine well-known natural products. Every correct LLM answer on these nine natural products names the molecule. **b**, Answers of Claude-Opus-5 on taxol and two taxanes. Of the other five taxanes, it identifies three in 1,178 to 1,480 s and reaches the output limit on two. GPT-5.6-Sol identifies four of the seven taxanes in 386 to 919 s and answers taxol in 732 s. Key sentences of the reasoning given with the first candidate are quoted, with omissions marked […]. The reasoning is the summary in the answer, not the model’s internal thinking.

| Molecule | Heavy atoms | LLM | Time (*s*) | First candidate | Confidence | Key sentences of the reasoning |
|---|---|---|---|---|---|---|
| **a** |  |  |  |  |  |  |
| Vancomycin aglycone | 80 | Claude-Opus-5 | 620 | correct | 0.78 | “C53H52Cl2N8O17 corresponds exactly to vancomycin aglycone (vancomycin C66H75Cl2N9O24 minus the glucose–vancosamine disaccharide, +H2O). [. . . ]” |
| Ginkgolide C | 31 | GPT-5.6-Sol | 1,081 | incorrect ring system | 0.99 | “The three lactone carbonyls, highly oxygenated hexacyclic cage, 9H tert-butyl singlet, secondary methyl doublet, and 18 carbon resonances (three equivalent tert-butyl methyls) identify ginkgolide C.” |
| Morphine | 21 | Gemini-3.6-Flash | 14 | incorrect, C<sub>16</sub>H<sub>19</sub>NO<sub>3</sub> instead of the given C<sub>17</sub>H<sub>19</sub>NO<sub>3</sub> | 0.95 | “The NMR spectral data matches morphine perfectly. [. . . ]” |
| **b** |  |  |  |  |  |  |
| Taxol | 62 | Claude-Opus-5 | 71 | correct | 0.95 | “*The molecular formula C47H51NO14 matches paclitaxel (Taxol) exactly.* The 1H pattern is diagnostic: [. . . ] 6.27 (s, H10), 6.23 (t, H13), 5.78 (dd, H3’) [. . . ]” |
| 1*β*-Hydroxytaxusin | 37 | Claude-Opus-5 | 880 | correct | 0.44 | “DoU 9; 8 methyl singlets (24H) = 4 OAc [. . . ] + 4 skeletal Me [. . . ] => taxane diterpene. [. . . ] No CH-OH signal, but a 5th oxygenated carbon at 77.0 plus OH at 1.42 s => tertiary 1*β*-OH [. . . ] Structure = 1*β*-hydroxytaxusin [. . . ]” |
| 4*α*,20-Epoxy-5*α*-hydroxytaxadien-13-one | 23 | Claude-Opus-5 | 1,514 | no answer (output limit) | – | none |

**Table S2.** Architecture and training settings of ChefNMR2. RMSNorm, root-mean-square normalization. SwiGLU, gated linear unit with sigmoid linear unit activation. MLP, multilayer perceptron. EMA, exponential moving average.

| Parameter | Value |
|---|---|
| **Peak encoder** |  |
| <sup>1</sup>H *peak embedding* |  |
| Number of Fourier frequencies *K* | 256 |
| Fourier frequencies *f<sub>k</sub>* | 1 to 64, logarithmically spaced |
| Shift range [*δ*<sub>min</sub>, *δ*<sub>max</sub>] | −2 to 12 ppm |
| Multiplicity | 128-dimensional learnable embedding, 8 classes |
| Number of protons | 128-dimensional learnable embedding, 0 to 20 |
| <sup>13</sup>C *peak embedding* |  |
| Number of Fourier frequencies *K* | 384 |
| Fourier frequencies *f<sub>k</sub>* | 1 to 128, logarithmically spaced |
| Shift range [*δ*<sub>min</sub>, *δ*<sub>max</sub>] | −20 to 230 ppm |
| *Transformer encoder* |  |
| Number of blocks | 4 |
| Hidden dimension | 768 |
| Number of attention heads | 8 |
| Feed-forward | SwiGLU, MLP ratio 4 |
| Normalization | RMSNorm, before each sublayer and on queries and keys |
| Positional encoding | None |
| Nucleus embedding | 768-dimensional learnable embedding, 2 classes |
| Dropout | 0.1 |
| *Pooling* |  |
| Pooling strategy | Multihead self-attention pooling |
| **Multimodal diffusion transformer** |  |
| Number of blocks | 12 |
| Hidden dimension | 768 |
| Number of attention heads | 8 |
| Feed-forward | SwiGLU, MLP ratio 4 |
| Normalization | RMSNorm, before each sublayer and on queries and keys |
| Atom positional encoding | None |
| **Training** |  |
| Trainable parameters | 288M |
| Optimizer | Adam (*β*<sub>1</sub> = 0.9, *β*<sub>2</sub> = 0.95, *ϵ* = 10<sup>−8</sup>) |
| Learning rate | 10<sup>−4</sup>, constant |
| Batch size | 1,024 |
| Epochs | 1,400 (3,452,400 training steps) |
| EMA decay | 0.999 |
| Precision | Mixed (bfloat16) |
| Dataset weights | 1 : 1 : 1 |
| Checkpoint ensemble | 10 checkpoints, at epochs 950, 1,000, …, 1,400 |

**Table S3.** Settings and prompts for the reasoning LLMs. Output limit, the largest number of output tokens per request. Price, the list price in US dollars per million input and output tokens, with reasoning tokens counted as output tokens. GLM-5.2, Kimi-K3 and DeepSeek-V4-Flash are accessed through OpenRouter, the others through their developers’ APIs. On the four novel natural products, Claude-Opus-5 is accessed through Anthropic’s Claude Code command-line interface, without tools or web access. Below, the system prompt and the user prompt template that every model receives for every molecule.

| System | Developer | Reasoning setting | Output limit | Price (in / out) |
|---|---|---|---|---|
| Claude-Opus-5 | Anthropic | effort max | 128,000 | 5.00 / 25.00 |
| GPT-5.6-Sol | OpenAI | effort max | 128,000 | 5.00 / 30.00 |
| Gemini-3.6-Flash | Google | thinking level high | 65,536 | 1.50 / 7.50 |
| GLM-5.2 | Z.ai | effort xhigh | 128,000 | 1.40 / 4.40 |
| Kimi-K3 | Moonshot AI | effort max | 128,000 | 3.00 / 15.00 |
| DeepSeek-V4-Flash | DeepSeek | effort xhigh | 128,000 | 0.14 / 0.28 |

**Table S4.** Molecular formulae and peak lists of the four novel natural products.

| Compound | Molecular formula | <sup>1</sup>H NMR *δ* | <sup>13</sup>C NMR *δ* |
|---|---|---|---|
| Nocobactin AI-1 | C<sub>38</sub>H<sub>61</sub>N<sub>5</sub>O<sub>11</sub> | 8.34 (m, 1H), 8.10 (d, 1H), 7.93 (m, 1H), 7.35 (q, 1H), 6.88 (td, 2H), 4.88 (td, 1H), 4.60 (q, 1H), 4.42 (ddd, 1H), 4.27 (tt, 1H), 3.86 (dd, 1H), 3.70 (d, 2H), 3.48 (m, 2H), 3.41 (t, 3H), 2.59 (m, 1H), 1.34 (m, 37H), 0.93 (d, 3H), 0.85 (t, 4H) | 172.2, 172.1, 172.0, 171.7, 171.7, 171.4, 169.9, 169.0, 167.7, 167.6, 163.6, 161.9, 161.6, 158.9, 158.8, 157.0, 133.5, 129.2, 118.8, 118.7, 117.8, 117.1, 116.4, 79.2, 75.8, 75.5, 61.7, 55.6, 52.3, 51.3, 50.7, 50.7, 49.0, 45.5, 43.7, 43.5, 32.0, 31.3, 31.3, 31.3, 30.6, 30.5, 30.1, 29.3, 29.1, 29.1, 29.1, 29.0, 29.0, 28.9, 28.9, 28.9, 28.8, 28.8, 28.7, 26.9, 26.6, 25.7, 24.9, 22.4, 22.2, 22.1, 22.1, 14.5, 14.5, 14.0, 14.0 |
| Nocobactin AI-2 | C<sub>36</sub>H<sub>55</sub>N<sub>5</sub>O<sub>10</sub> | 11.79 (s, 1H), 9.75 (s, 2H), 8.73 (dd, 1H), 8.39 (s, 1H), 8.21 (s, 1H), 8.11 (dd, 1H), 7.86 (s, 1H), 7.63 (dd, 1H), 7.46 (ddd, 1H), 7.00 (d, 1H), 6.94 (t, 1H), 5.01 (dd, 1H), 4.89 (td, 1H), 4.65 (dd, 1H), 4.45 (m, 2H), 4.25 (m, 1H), 3.86 (ddd, 1H), 3.46 (m, 2H), 3.37 (s, 1H), 2.61 (ddd, 1H), 1.77 (m, 2H), 1.67 (dd, 2H), 1.62 (m, 1H), 1.54 (q, 2H), 1.43 (tt, 3H), 1.34 (m, 2H), 1.22 (m, 6H), 1.19 (s, 10H), 1.10 (m, 2H), 0.94 (dd, 4H), 0.84 (t, 4H) | 172.2, 171.6, 170.0, 169.0, 166.0, 165.8, 161.6, 159.1, 157.0, 134.1, 128.1, 119.1, 116.6, 109.9, 75.7, 69.3, 67.1, 52.7, 52.3, 50.7, 49.0, 45.4, 43.7, 40.4, 32.0, 31.3, 30.3, 30.3, 30.2, 29.0, 29.0, 28.9, 28.9, 28.8, 28.8, 28.7, 27.0, 26.5, 25.7, 24.8, 22.6, 22.4, 22.2, 14.6, 14.0 |
| Hybridamide A | C<sub>25</sub>H<sub>36</sub>N<sub>6</sub>O<sub>4</sub> | 7.55 (dt, 1H), 7.32 (dd, 1H), 7.07 (m, 2H), 7.00 (ddd, 1H), 5.14 (tp, 1H), 4.34 (dd, 1H), 3.92 (d, 2H), 3.46 (t, 2H), 3.32 (d, 1H), 3.25 (dt, 1H), 2.93 (m, 2H), 2.49 (m, 4H), 1.82 (dq, 1H), 1.75 (d, 3H), 1.69 (m, 6H) | 176.8, 174.5, 174.4, 166.6, 157.7, 139.6, 138.2, 128.8, 123.5, 122.3, 119.6, 119.3, 118.7, 113.3, 112.2, 54.1, 48.7, 47.4, 41.5, 32.3, 32.2, 30.5, 26.3, 25.8, 24.3, 18.0 |
| Hybridamide B | C<sub>20</sub>H<sub>28</sub>N<sub>6</sub>O<sub>4</sub> | 7.55 (d, 1H), 7.31 (m, 2H), 7.07 (m, 2H), 7.00 (ddd, 1H), 4.38 (dd, 1H), 3.46 (t, 2H), 3.18 (hept, 2H), 2.93 (t, 2H), 2.50 (m, 4H), 1.91 (m, 1H), 1.67 (m, 3H) | 176.1, 174.6, 158.6, 138.2, 128.8, 123.4, 122.3, 119.6, 119.2, 113.3, 112.2, 53.7, 42.0, 41.5, 32.2, 32.1, 30.3, 26.2, 26.1 |

**Algorithm S1 | Smooth lDDT loss.**

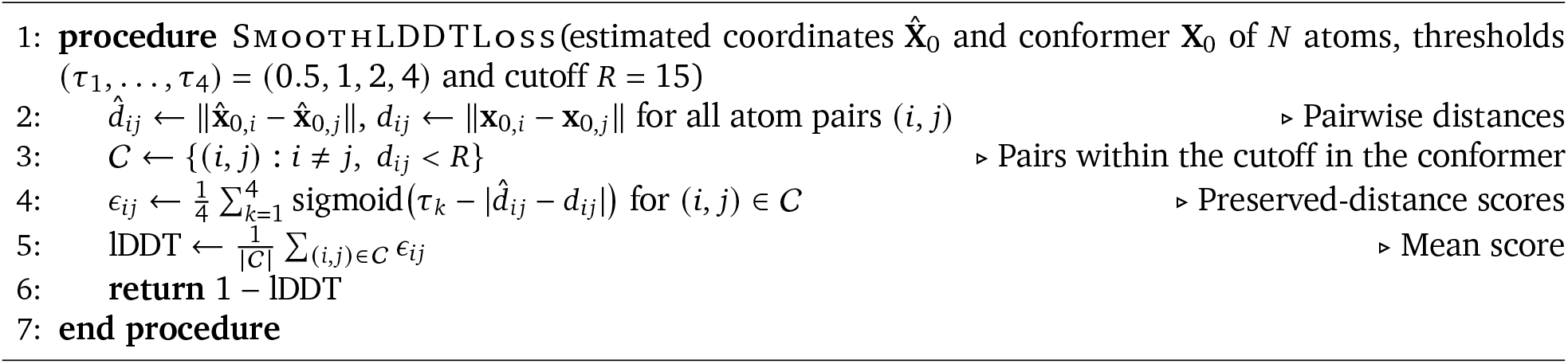

**Algorithm S2 | Training of ChefNMR2.**

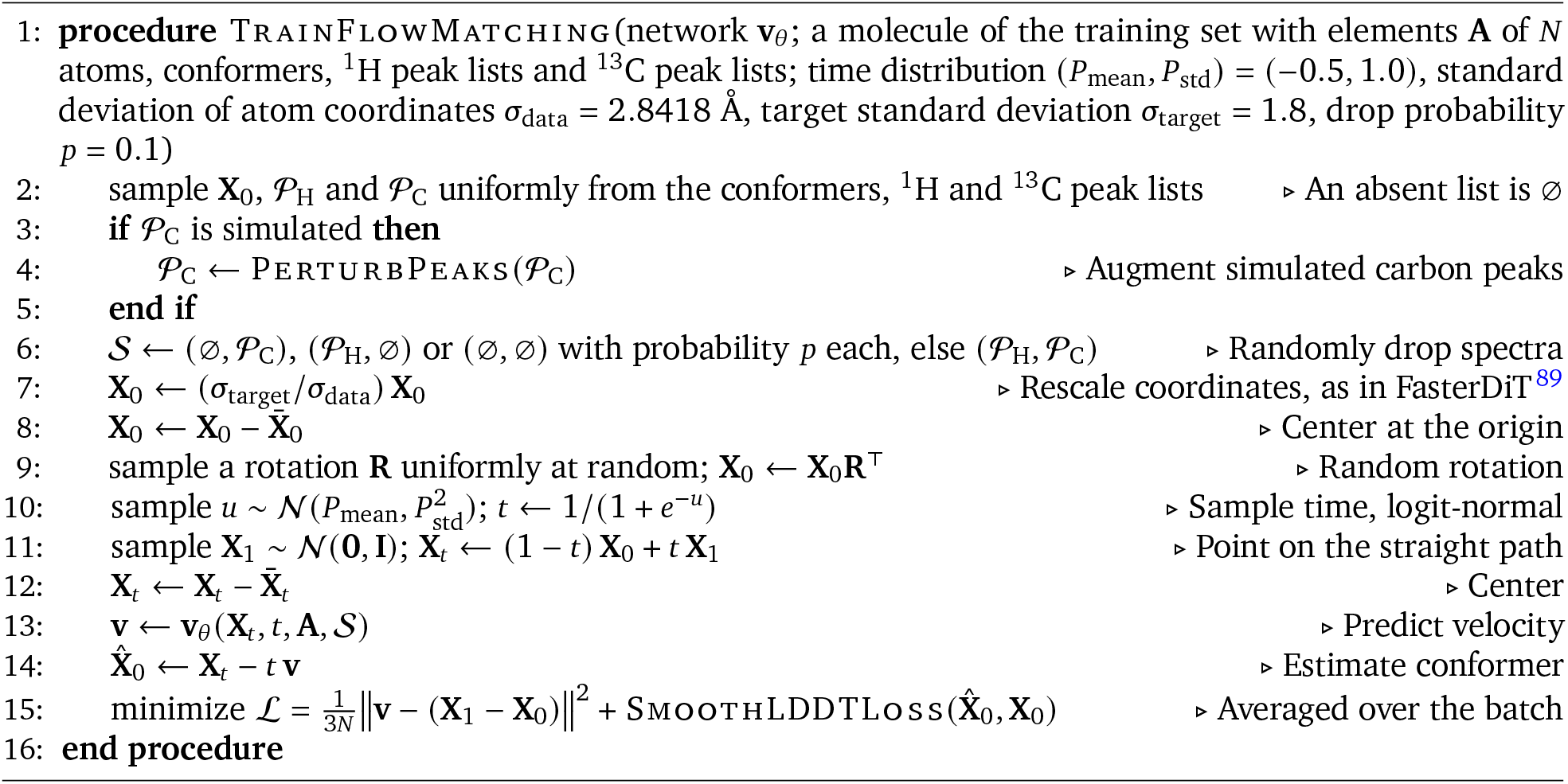

**Algorithm S3 | Generation of one sample by ChefNMR2.**

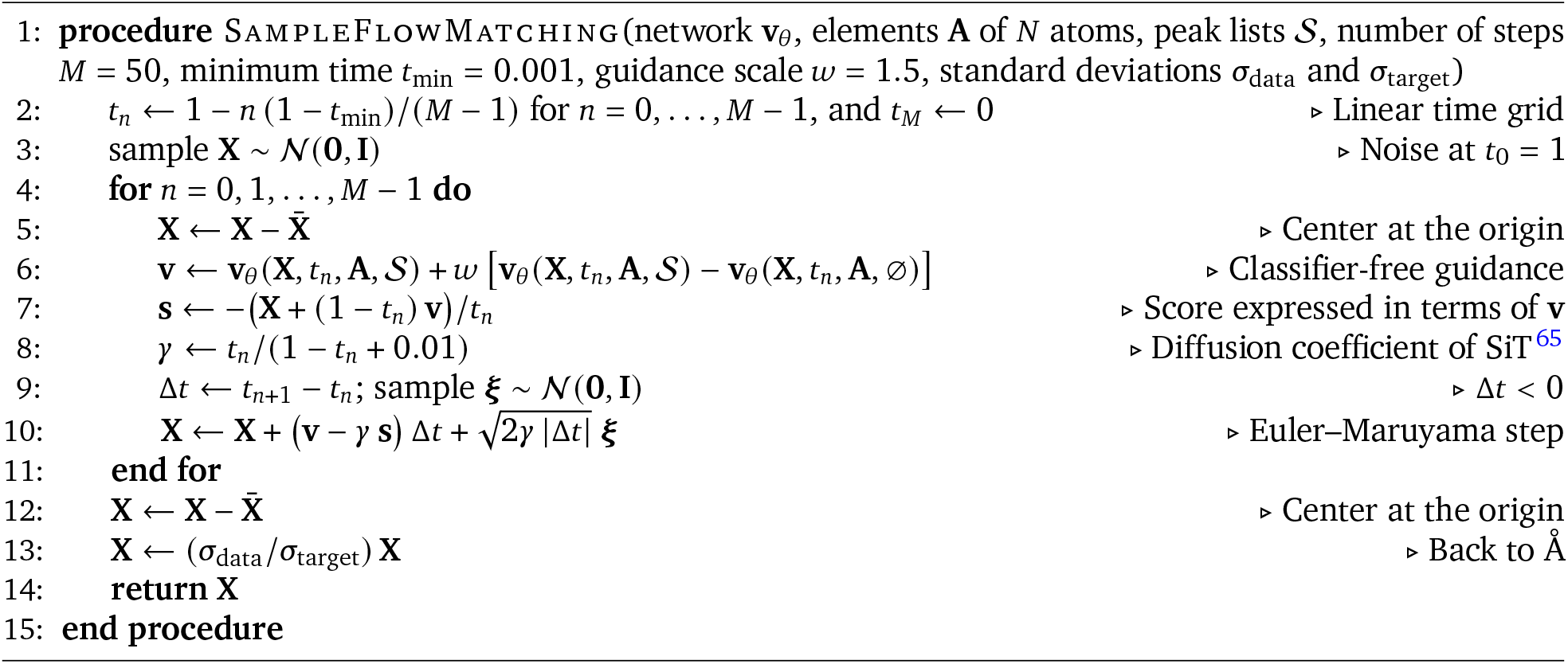

**Algorithm S4 | Perturbation of a simulated ^13^C peak list.**

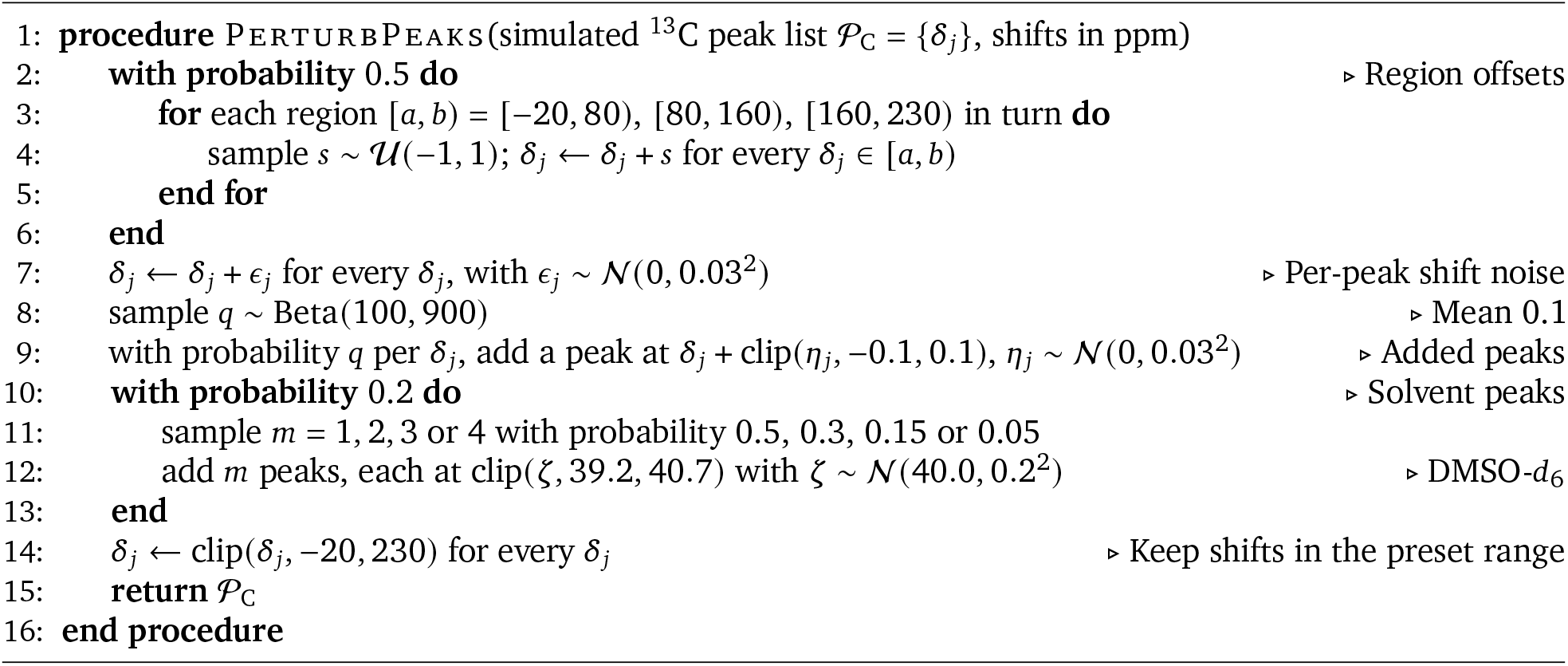

**Algorithm S5 | Adaptive sampling and ranking of structures by ChefNMR2.**

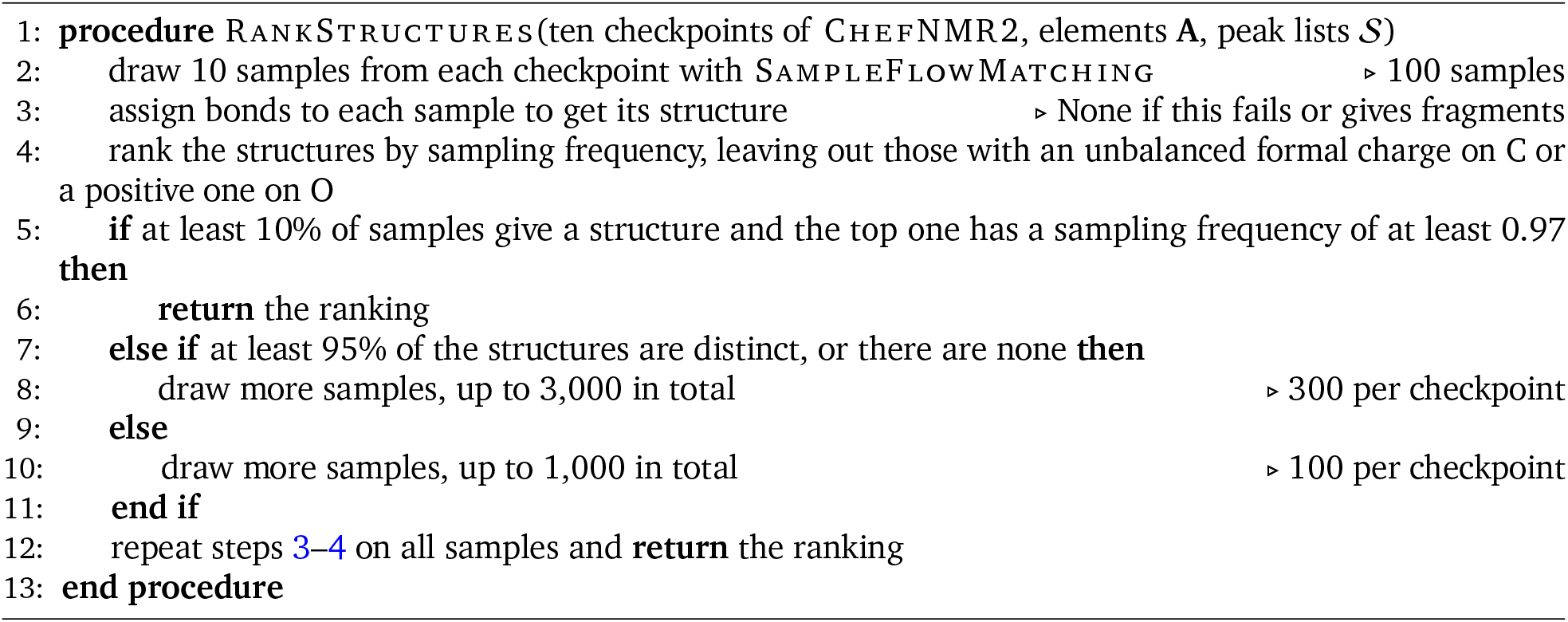

## References

[1] David J Newman and Gordon M Cragg. Natural products as sources of new drugs over the nearly four decades from 01/1981 to 09/2019. Journal of Natural Products, 83(3):770–803, 2020.

[2] Jon Clardy and Christopher Walsh. Lessons from natural molecules. Nature, 432(7019):829–837, 2004.

[3] Gina Porras, François Chassagne, James T Lyles, Lewis Marquez, Micah Dettweiler, Akram M Salam, Tharanga Samarakoon, Sarah Shabih, Darya Raschid Farrokhi, and Cassandra L Quave. Ethnobotany and the role of plant natural products in antibiotic drug discovery. Chemical Reviews, 121(6):3495–3560, 2021.

[4] Yizhou Liu, Josep Saurí, Emily Mevers, Mark W Peczuh, Henk Hiemstra, Jon Clardy, Gary E Martin, and R Thomas Williamson. Unequivocal determination of complex molecular structures using anisotropic NMR measurements. Science, 356(6333):eaam5349, 2017.

[5] Ēriks Kupče, Lucio Frydman, Andrew G Webb, Jonathan R J Yong, and Tim D W Claridge. Parallel nuclear magnetic resonance spectroscopy. Nature Reviews Methods Primers, 1(1):27, 2021.

[6] Frank Hu, Michael S Chen, Grant M Rotskoff, Matthew W Kanan, and Thomas E Markland. Accurate and efficient structure elucidation from routine one-dimensional NMR spectra using multitask machine learning. ACS Central Science, 10(11):2162–2170, 2024.

[7] Mikhail Elyashberg, Kirill Blinov, Sergey Molodtsov, Yegor Smurnyy, Antony J Williams, and Tatiana Churanova. Computer-assisted methods for molecular structure elucidation: realizing a spectroscopist’s dream. Journal of Cheminformatics, 1(1):3, 2009.

[8] Darcy C Burns, Eugene P Mazzola, and William F Reynolds. The role of computer-assisted structure elucidation (CASE) programs in the structure elucidation of complex natural products. Natural Product Reports, 36(6):919–933, 2019.

[9] Lucio Frydman, Tali Scherf, and Adonis Lupulescu. The acquisition of multidimensional NMR spectra within a single scan. Proceedings of the National Academy of Sciences, 99(25):15858– 15862, 2002.

[10] Frank Hu, Jonathan M Tubb, Dimitris Argyropoulos, Sergey Golotvin, Mikhail Elyashberg, Grant M Rotskoff, Matthew W Kanan, and Thomas E Markland. Pushing the limits of one-dimensional NMR spectroscopy for automated structure elucidation using artificial intelligence. Journal of Chemical Information and Modeling, 66(12):6992–7002, 2026.

[11] Marvin Alberts, Nina Hartrampf, and Teodoro Laino. Automated structure elucidation at human-level accuracy via a multimodal multitask language model. ChemRxiv, 2025.

[12] Xi Xue, Hanyu Sun, Jingying Sun, Luc Patiny, Xiangying Liu, Kai Chen, Jingjie Yan, Liangning Li, Xue Liu, Shu Xu, Dongming Zhang, Yafeng Deng, Yingda Zang, Yaling Gong, Jie Ma, and Xiaojian Wang. NMRMind: a transformer-based model enabling the elucidation from multidimensional NMR to structures. Analytical Chemistry, 97(41):22603–22614, 2025.

[13] Fanjie Xu, Jinyuan Hu, Jingxiang Zou, Junjie Wang, Boying Huang, Zhifeng Gao, Xiaohong Ji, Weinan E, Zhong-Qun Tian, Fujie Tang, and Jun Cheng. NMRPeak: a ready-to-use intelligent system for molecular structure elucidation enabled by synergistic cross-modal learning. arXiv preprint arXiv:2602.08752, 2026.

[14] Liujia Yang, Zhuo Yang, Jiaqing Xie, Yubin Wang, Ben Gao, Xingjian Wei, Jiaxing Sun, Jiang Wu, Conghui He, Yuqiang Li, and Qinying Gu. NMRTrans: structure elucidation from experimental NMR spectra via set transformers. In Proceedings of the 32nd ACM SIGKDD Conference on Knowledge Discovery and Data Mining V.2, pages 12621–12632, 2026.

[15] Chen Yang, Zheng Fang, Hanyu Sun, Fanjie Xu, Hongxin Xiang, Hanyu Gao, Xiangxiang Zeng, Yuqiang Li, Xiaojian Wang, and Jun Xia. A large-scale foundation model enables simulation-to-real adaptation for nuclear magnetic resonance-based molecular structure analysis. arXiv preprint arXiv:2606.20756, 2026.

[16] Bingsen Xue, Zhuojun Jiang, Jianhao Zhang, Mingcheng Gu, Yizhe Yuan, Yongtai Zhuo, Yifan Zhang, Li Wang, Ya Su, Yue Yuan, Jiang Liu, Xueqian Kong, and Cheng Jin. Multi-agent closed-loop reasoning for organic structure elucidation from multimodal spectra. arXiv preprint arXiv:2608.14720, 2026.

[17] Ivan Zlobin, Ivan Bespalov, Maksim Shandybo, Nikolai Rekut, Yulia Nelyubina, Denis Chusov, Oleg Afanasyev, and Valentin Novikov. Bimodal learning from imperfect NMR data (BLIND): a chemistry-ignorant model for autonomous structure elucidation from NMR. ChemRxiv, 2026.

[18] Ziyu Xiong, Yichi Zhang, Foyez Alauddin, Chu Xin Cheng, Joon Soo An, Mohammad R Seyedsayamdost, and Ellen D Zhong. Atomic diffusion models for small molecule structure elucidation from NMR spectra. In Advances in Neural Information Processing Systems, volume 38, pages 128593–128629, 2025.

[19] Yongqi Jin, Jun-Jie Wang, Fanjie Xu, Xiaohong Ji, Zhifeng Gao, Linfeng Zhang, Guolin Ke, Rong Zhu, and Weinan E. NMR-Solver: automated structure elucidation via large-scale spectral matching and physics-guided fragment optimization. Nature Communications, 17(1):4740, 2026.

[20] Adrian Mirza, Luc Patiny, and Kevin Maik Jablonka. End-to-end multimodal structure elucidation from raw spectra combining contrastive learning and evolutionary algorithms. Nature Communications, 17(1):5013, 2026.

[21] Marvin Alberts, Oliver Schilter, Federico Zipoli, Nina Hartrampf, and Teodoro Laino. Unraveling molecular structure: A multimodal spectroscopic dataset for chemistry. In Advances in Neural Information Processing Systems, volume 37, pages 125780–125808, 2024.

[22] Zihan Zhang, Mengxi Chen, Xuezhou Zhao, Yutao Guo, and Dan Wu. Public database leak-age distorts model rankings in real-spectra NMR structure elucidation. Journal of Chemical Information and Modeling, 66(19):12637–12644, 2026.

[23] Siddharth Narayanan, James Braza, Ryan-Rhys Griffiths, Albert Bou, Geemi Wellawatte, Mayk Caldas Ramos, Ludovico Mitchener, Michael Pieler, Sam Rodriques, and Andrew White. Training a scientific reasoning model for chemistry. In Advances in Neural Information Processing Systems, volume 38, pages 175068–175107, 2025.

[24] Nicholas T Runcie, Charlotte M Deane, and Fergus Imrie. Assessing the chemical intelligence of large language models. Journal of Chemical Information and Modeling, 66(1):216–227, 2026.

[25] Samiha Sharlin, Fariha Agbere, Kevin Ishimwe, Zuzana Osifová, Ondřej Socha, Martin Dračínský, and Tyler R Josephson. NMR-Challenge for LLMs: evaluating chemical reasoning in humans and AI. Journal of Chemical Information and Modeling, 66(11):6299–6312, 2026.

[26] Fedor Kliuev, Ivan Smirnov, Mikhail A Losev, Oleg I Afanasyev, and Denis Chusov. Can AI help chemists to solve NMR? ChemRxiv, 2026.

[27] Zheng Fang, Chen Yang, Yusen Tan, Yunpeng Zhao, Fanjie Xu, Hongxin Xiang, Hanyu Sun, Hanyu Gao, Xiaojian Wang, Wenjie Du, Yuqiang Li, and Jun Xia. Towards generalizable and evidential nuclear magnetic resonance-based molecular structure elucidation via large language model agent. arXiv preprint arXiv:2606.29776, 2026.

[28] Daniil A Boiko, Robert MacKnight, Ben Kline, and Gabe Gomes. Autonomous chemical research with large language models. Nature, 624(7992):570–578, 2023.

[29] Andres M Bran, Sam Cox, Oliver Schilter, Carlo Baldassari, Andrew D White, and Philippe Schwaller. Augmenting large language models with chemistry tools. Nature Machine Intelligence, 6(5):525–535, 2024.

[30] Ludovico Mitchener, Angela Yiu, Benjamin Chang, Mathieu Bourdenx, Tyler Nadolski, Arvis Sulovari, Eric C Landsness, Daniel L Barabasi, Siddharth Narayanan, Nicky Evans, Shriya Reddy, Martha Foiani, Aizad Kamal, Leah P Shriver, Fang Cao, Asmamaw T Wassie, Jon M Laurent, Edwin Melville-Green, Mayk Caldas, Albert Bou, Kaleigh F Roberts, Sladjana Zagorac, Timothy C Orr, Miranda E Orr, Kevin J Zwezdaryk, Ali E Ghareeb, Laurie McCoy, Bruna Gomes, Euan A Ashley, Karen E Duff, Tonio Buonassisi, Tom Rainforth, Randall J Bateman, Michael Skarlinski, Samuel G Rodriques, Michaela M Hinks, and Andrew D White. Kosmos: an AI scientist for autonomous discovery. arXiv preprint arXiv:2511.02824, 2025.

[31] Martin Priessner, Richard J Lewis, Magnus J Johansson, Jonathan M Goodman, Jon Paul Janet, and Anna Tomberg. Enhancing molecular structure elucidation with reasoning-capable LLMs. Digital Discovery, 5(3):1237–1251, 2026.

[32] Yutang Ge, Yaning Cui, Hanzheng Li, Jun-Jie Wang, Fanjie Xu, Jinhan Dong, Yongqi Jin, Dongxu Cui, Peng Jin, Guojiang Zhao, Hengxing Cai, Tianci Yangfeng, Xueqing Chen, Hongshuai Wang, Rong Zhu, Linfeng Zhang, Xiaohong Ji, and Zhifeng Gao. SpecXMaster technical report. arXiv preprint arXiv:2603.23101, 2026.

[33] Patrick Esser, Sumith Kulal, Andreas Blattmann, Rahim Entezari, Jonas Müller, Harry Saini, Yam Levi, Dominik Lorenz, Axel Sauer, Frederic Boesel, Dustin Podell, Tim Dockhorn, Zion English, Kyle Lacey, Alex Goodwin, Yannik Marek, and Robin Rombach. Scaling rectified flow transformers for high-resolution image synthesis. In International Conference on Machine Learning, 2024.

[34] Yaron Lipman, Ricky T. Q. Chen, Heli Ben-Hamu, Maximilian Nickel, and Matt Le. Flow matching for generative modeling. In International Conference on Learning Representations, 2023.

[35] Josh Abramson, Jonas Adler, Jack Dunger, Richard Evans, Tim Green, Alexander Pritzel, Olaf Ronneberger, Lindsay Willmore, Andrew J Ballard, Joshua Bambrick, et al. Accurate structure prediction of biomolecular interactions with AlphaFold 3. Nature, 630(8016):493–500, 2024.

[36] Jun-Jie Wang, Yongqi Jin, Chen-Yu Zhi, Yu-Jie Liu, Xu-Hao Huang, Fanjie Xu, Xiaohong Ji, Xi Fang, Haoyi Tao, Weinan E, Linfeng Zhang, Guolin Ke, and Rong Zhu. NMRexp: A database of 3.3 million experimental NMR spectra. Scientific Data, 12(1):1954, 2025.

[37] Daniel Mark Lowe. Extraction of chemical structures and reactions from the literature. PhD thesis, University of Cambridge, 2012.

[38] Venkata Chandrasekhar, Kohulan Rajan, Sri Ram Sagar Kanakam, Nisha Sharma, Viktor Weißenborn, Jonas Schaub, and Christoph Steinbeck. COCONUT 2.0: a comprehensive overhaul and curation of the collection of open natural products database. Nucleic Acids Research, 53(D1): D634–D643, 2025.

[39] Ella F Poynton, Jeffrey A van Santen, Matthew Pin, Marla Macias Contreras, Emily McMann, Jonathan Parra, Brandon Showalter, Liana Zaroubi, Katherine R Duncan, and Roger G Linington. The Natural Products Atlas 3.0: extending the database of microbially derived natural products. Nucleic Acids Research, 53(D1):D691–D699, 2025.

[40] David S Wishart, Tanvir Sajed, Matthew Pin, Ella F Poynton, Bharat Goel, Brian L Lee, An Chi Guo, Sukanta Saha, Zinat Sayeeda, Scott Han, et al. The Natural Products Magnetic Resonance Database (NP-MRD) for 2025. Nucleic Acids Research, 53(D1):D700–D708, 2025.

[41] J. Benji Rowlands, Richard J. Lewis, Peter Howe, and Jonathan M. Goodman. CSP5: Large-scale neural chemical shift prediction from 2.5 million experimental NMR spectra. ChemRxiv, 2026.

[42] Christoph Steinbeck and Stefan Kuhn. NMRShiftDB – compound identification and structure elucidation support through a free community-built web database. Phytochemistry, 65(19): 2711–2717, 2004.

[43] Stefan Kuhn and Nils E Schlörer. Facilitating quality control for spectra assignments of small organic molecules: nmrshiftdb2 – a free in-house NMR database with integrated LIMS for academic service laboratories. Magnetic Resonance in Chemistry, 53(8):582–589, 2015.

[44] Scott E Van Bramer and Loyd D Bastin. Spectroscopy data for undergraduate teaching. Journal of Chemical Education, 100(10):3897–3902, 2023.

[45] Mahdi Pakdaman Naeini, Gregory Cooper, and Milos Hauskrecht. Obtaining well calibrated probabilities using Bayesian binning. In Proceedings of the AAAI Conference on Artificial Intelligence, volume 29, 2015.

[46] Conor James McClune, Jack Chun-Ting Liu, Chloe Wick, Ricardo De La Peña, Bernd Markus Lange, Polly M. Fordyce, and Elizabeth S. Sattely. Discovery of FoTO1 and Taxol genes enables biosynthesis of baccatin III. Nature, 643(8071):582–592, 2025.

[47] Luis E N Quadri, Jason Sello, Thomas A Keating, Paul H Weinreb, and Christopher T Walsh. Identification of a Mycobacterium tuberculosis gene cluster encoding the biosynthetic enzymes for assembly of the virulence-conferring siderophore mycobactin. Chemistry & Biology, 5(11): 631–645, 1998.

[48] H Drechsel and G Jung. Peptide siderophores. Journal of Peptide Science, 4(3):147–181, 1998.

[49] Fabián Garzón-Posse, Yovanny Quevedo-Acosta, Camilo Mahecha-Mahecha, and Paola Acosta Guzmán. Recent progress in the synthesis of naturally occurring siderophores. European Journal of Organic Chemistry, 2019(48):7747–7769, 2019.

[50] Yunqiu Chen, Michelle Unger, Ioanna Ntai, Ryan A. McClure, Jessica C. Albright, Regan J. Thomson, and Neil L. Kelleher. Gobichelin A and B: mixed-ligand siderophores discovered using proteomics. MedChemComm, 4(1):233–238, 2013.

[51] Midori A Arai, Itsuki Ebihara, Yoshinori Makita, Yasumasa Hara, Takashi Yaguchi, and Masami Ishibashi. Isolation of nocobactin NAs as Notch signal inhibitors from Nocardia farcinica, a possibility of invasive evolution. The Journal of Antibiotics, 74(4):255–259, 2021.

[52] Z J Huang, Y V Ying, Z Zheng, J S An, E A Creasey, L M Taylor, D B Graham, R J Xavier, and M R Seyedsayamdost. Under revision, 2026.

[53] Sereina Riniker and Gregory A Landrum. Better informed distance geometry: using what we know to improve conformation generation. Journal of Chemical Information and Modeling, 55 (12):2562–2574, 2015.

[54] Shuzhe Wang, Jagna Witek, Gregory A Landrum, and Sereina Riniker. Improving conformer generation for small rings and macrocycles based on distance geometry and experimental torsional-angle preferences. Journal of Chemical Information and Modeling, 60(4):2044–2058, 2020.

[55] Greg Landrum, Paolo Tosco, Brian Kelley, et al. rdkit/rdkit: 2025_09_4 (Q3 2025) release. Zenodo, 2025.

[56] Thomas A Halgren. Merck molecular force field. I. Basis, form, scope, parameterization, and performance of MMFF94. Journal of Computational Chemistry, 17(5-6):490–519, 1996.

[57] Anthony K Rappé, Carla J Casewit, Kent S Colwell, William A Goddard III, and W Mason Skiff. UFF, a full periodic table force field for molecular mechanics and molecular dynamics simulations. Journal of the American Chemical Society, 114(25):10024–10035, 1992.

[58] Yeonjoon Kim and Woo Youn Kim. Universal structure conversion method for organic molecules: from atomic connectivity to three-dimensional geometry. Bulletin of the Korean Chemical Society, 36(7):1769–1777, 2015.

[59] Jonathan Bisson, Charlotte Simmler, Shao-Nong Chen, J Brent Friesen, David C Lankin, James B McAlpine, and Guido F Pauli. Dissemination of original NMR data enhances reproducibility and integrity in chemical research. Natural Product Reports, 33(9):1028–1033, 2016.

[60] Mestrelab Research. Mnova: NMR data processing and analysis software. https://mestrelab.com/main-product/mnova, 2025.

[61] David Weininger. SMILES, a chemical language and information system. 1. Introduction to methodology and encoding rules. Journal of Chemical Information and Computer Sciences, 28 (1):31–36, 1988.

[62] Nadine Schneider, Roger A. Sayle, and Gregory A. Landrum. Get your atoms in order—an open-source implementation of a novel and robust molecular canonicalization algorithm. Journal of Chemical Information and Modeling, 55(10):2111–2120, 2015.

[63] Xingchao Liu, Chengyue Gong, and Qiang Liu. Flow straight and fast: Learning to generate and transfer data with rectified flow. In International Conference on Learning Representations, 2023.

[64] Valerio Mariani, Marco Biasini, Alessandro Barbato, and Torsten Schwede. lDDT: a local superposition-free score for comparing protein structures and models using distance difference tests. Bioinformatics, 29(21):2722–2728, 2013.

[65] Nanye Ma, Mark Goldstein, Michael S. Albergo, Nicholas M. Boffi, Eric Vanden-Eijnden, and Saining Xie. SiT: Exploring flow and diffusion-based generative models with scalable interpolant transformers. In Computer Vision – ECCV 2024, Lecture Notes in Computer Science, pages 23–40, 2024.

[66] Jonathan Ho and Tim Salimans. Classifier-free diffusion guidance. arXiv preprint arXiv:2207.12598, 2022.

[67] William Peebles and Saining Xie. Scalable diffusion models with transformers. In 2023 IEEE/CVF International Conference on Computer Vision (ICCV), pages 4172–4182, 2023.

[68] Matthew Tancik, Pratul Srinivasan, Ben Mildenhall, Sara Fridovich-Keil, Nithin Raghavan, Utkarsh Singhal, Ravi Ramamoorthi, Jonathan Barron, and Ren Ng. Fourier features let networks learn high frequency functions in low dimensional domains. Advances in Neural Information Processing Systems, 33:7537–7547, 2020.

[69] Juho Lee, Yoonho Lee, Jungtaek Kim, Adam Kosiorek, Seungjin Choi, and Yee Whye Teh. Set transformer: A framework for attention-based permutation-invariant neural networks. In International Conference on Machine Learning, pages 3744–3753. PMLR, 2019.

[70] Hugo Touvron, Matthieu Cord, Alexandre Sablayrolles, Gabriel Synnaeve, and Hervé Jégou. Going deeper with image transformers. In 2021 IEEE/CVF International Conference on Computer Vision (ICCV), pages 32–42. IEEE, 2021.

[71] Chaitanya K Joshi, Xiang Fu, Yi-Lun Liao, Vahe Gharakhanyan, Benjamin Kurt Miller, Anuroop Sriram, and Zachary W Ulissi. All-atom Diffusion Transformers: Unified generative modelling of molecules and materials. In International Conference on Machine Learning, 2025.

[72] Zhiyuan Liu, Yanchen Luo, Han Huang, Enzhi Zhang, Sihang Li, Junfeng Fang, Yaorui Shi, Xiang Wang, Kenji Kawaguchi, and Tat-Seng Chua. NExT-Mol: 3D diffusion meets 1D language modeling for 3D molecule generation. In International Conference on Learning Representations, 2025.

[73] Diederik P Kingma and Jimmy Ba. Adam: A method for stochastic optimization. arXiv preprint arXiv:1412.6980, 2014.

[74] Joël Turconi, Frédéric Griolet, Ronan Guevel, Gilles Oddon, Roberto Villa, Andrea Geatti, Massimo Hvala, Kai Rossen, Rudolf Göller, and Andreas Burgard. Semisynthetic artemisinin, the chemical path to industrial production. Organic Process Research & Development, 18(3): 417–422, 2014.

[75] Martin Hébert, Gabriel Bellavance, and Louis Barriault. Total synthesis of ginkgolide C and formal syntheses of ginkgolides A and B. Journal of the American Chemical Society, 144(39): 17792–17796, 2022.

[76] Lars Jørgensen, Steven J. McKerrall, Christian A. Kuttruff, Felix Ungeheuer, Jakob Felding, and Phil S. Baran. 14-step synthesis of (+)-ingenol from (+)-3-carene. Science, 341(6148):878–882, 2013.

[77] Qiang Liu, Yu-An Zhang, Ping Xu, and Yanxing Jia. Total synthesis of (+)-lysergic acid. The Journal of Organic Chemistry, 78(21):10885–10893, 2013.

[78] Shuyu Chu, Niels Münster, Tudor Balan, and Martin D. Smith. A cascade strategy enables a total synthesis of (±)-morphine. Angewandte Chemie International Edition, 55(46):14306–14309, 2016.

[79] James B. McAlpine, Susan J. Swanson, Marianna Jackson, and David N. Whittern. Revised NMR assignments for rapamycin. The Journal of Antibiotics, 44(6):688–690, 1991.

[80] K. C. Nicolaou, P. G. Nantermet, H. Ueno, R. K. Guy, E. A. Couladouros, and E. J. Sorensen. Total synthesis of Taxol. 1. retrosynthesis, degradation, and reconstitution. Journal of the American Chemical Society, 117(2):624–633, 1995.

[81] Christopher Thompson, Min Ge, and Daniel Kahne. Synthesis of vancomycin from the aglycon. Journal of the American Chemical Societ*y*, 121(6):1237–1244, 1999.

[82] Tero Karras, Miika Aittala, Timo Aila, and Samuli Laine. Elucidating the design space of diffusion-based generative models. Advances in neural information processing systems, 35:26565–26577, 2022.

[83] Fanjie Xu, Wentao Guo, Feng Wang, Lin Yao, Hongshuai Wang, Fujie Tang, Zhifeng Gao, Linfeng Zhang, Weinan E, Zhong-Qun Tian, and Jun Cheng. Toward a unified benchmark and framework for deep learning-based prediction of nuclear magnetic resonance chemical shifts. Nature Computational Science, 5(4):292–300, 2025.

[84] Dávid Bajusz, Anita Rácz, and Károly Héberger. Why is Tanimoto index an appropriate choice for fingerprint-based similarity calculations? Journal of Cheminformatics, 7(1):20, 2015.

[85] David Rogers and Mathew Hahn. Extended-connectivity fingerprints. Journal of Chemical Information and Modeling, 50(5):742–754, 2010.

[86] Modal Labs. Pricing. https://modal.com/pricing, 2026. Accessed 28 September 2026.

[87] Sunghwan Kim, Jie Chen, Tiejun Cheng, Asta Gindulyte, Jia He, Siqian He, Qingliang Li, Benjamin A Shoemaker, Paul A Thiessen, Bo Yu, Leonid Zaslavsky, Jian Zhang, and Evan E Bolton. PubChem 2025 update. Nucleic Acids Research, 53(D1):D1516–D1525, 2025.

[88] Leland McInnes, John Healy, and James Melville. UMAP: Uniform manifold approximation and projection for dimension reduction. arXiv preprint arXiv:1802.03426, 2018.

[89] Jingfeng Yao, Cheng Wang, Wenyu Liu, and Xinggang Wang. FasterDiT: Towards faster diffusion transformers training without architecture modification. Advances in Neural Information Processing Systems, 37:56166–56189, 2024.

